# An atlas of cell type specific regulatory effects in cattle

**DOI:** 10.1101/2025.06.23.661035

**Authors:** Houcheng Li, Huicong Zhang, Pengju Zhao, Qi Zhang, Senlin Zhu, Tao Shi, Ya-Nan Wang, Jing-Sheng Lu, Liu Yang, John F. O’Grady, David E. MacHugh, Yu Wang, Zhenyu Wei, Xuemei Lu, Ming-Shan Wang, Bo Han, Weijie Zheng, Ao Chen, Shamima Akter, Nayan Bhowmik, Ying Ma, Ransom L. Baldwin, Congjun Li, Jicai Jiang, Li Ma, Christian Maltecca, Junjian Wang, Mian Gong, Xiaoning Zhu, Qing Lin, Yang Xi, Di Zhu, Jinyan Teng, Dailu Guan, Bingxing An, Jilong Ren, Yali Hou, Fei Wang, Bingjie Li, Laurent A. F. Frantz, Greger Larson, Zexi Cai, Goutam Sahana, Yu Jiang, Huizeng Sun, Dongxiao Sun, Geroge E. Liu, Lingzhao Fang

## Abstract

Understanding the genetic and molecular architecture of complex traits and artificial selection is crucial for advancing sustainable precision breeding in cattle and other livestock. Yet, how genetic variation affects cellular gene expression remains elusive in cattle. Here, by integrating 8,866 bulk RNA-seq samples and 999,192 single cells of 81 cell types in 22 bovine tissues, we presented a comprehensive atlas of regulatory variants at the cell type resolution in cattle. By colocalizing with bulk-tissue expression quantitative trait loci (beQTL), we detected 57,043 novel cell-type stratified eQTL and cell-type/state interaction eQTL in 18,153 genes, which also exhibited a stronger tissue/cell-type specificity than beQTL. By examining genome-wide associations (GWAS) of 44 complex traits, these cell-resolved eQTL were colocalized with 505 (24%) additional GWAS loci compared to beQTL. Through integrating this resource with selection signatures between dairy and beef cattle, we provided tissue/cell-specific regulatory insights into cattle breeding. Overall, the current atlas of cell-type-specific regulatory variants will serve as an invaluable resource for cattle genomics and selective breeding.

## Introduction

Cattle (*Bos taurus*), domesticated over 10,000 years ago, are among the most economically important livestock species worldwide^1^. They efficiently convert indigestible plant fibers into high-quality, protein-rich products such as milk and beef, which are important for human nutrition and health^2,3^. Over millennia, and particularly through recent intensive breeding, cattle have undergone extensive human-mediated selection to improve productivity, fertility, disease resistance, and adaptability^4^. Through long-term adaptation within a wide range of environmental milieus, cattle have further diversified into multiple breeds with distinct physiological and morphological characteristics^5,6^. In the face of rising global demand for sustainable and safe animal-derived food products, alongside growing concerns over environmental impacts (e.g., greenhouse gas emissions) and animal welfare^7,8^, it is critical to understand the genetic and molecular mechanisms that underlie economically and ecologically important traits in cattle.

Previous efforts, such as those led by the Farm Animal Genotype-Tissue Expression (FarmGTEx) and the Functional Annotation of Animal Genomes (FAANG) consortia, have made significant advances in characterizing the functional and regulatory genomes of a variety of farmed animal species (e.g., cattle, pigs, and chickens)^9–12^. These studies primarily leveraged bulk tissue RNA-sequencing (RNA-seq) data to integrate and associate molecular phenotype variation with complex trait variation. For instance, the CattleGTEx described the transcriptomic landscape of 23 distinct tissues and linked genomic variation impacting gene expression in these tissues to complex traits in cattle^9^. However, the inherent heterogeneity of bulk tissues—comprising diverse cell types and states—limits the cellular resolution at which regulatory mechanisms can be resolved. As genetic variants often exert their regulatory effects in a cell-type or cell-state specific manner^13–15^, deciphering their function requires knowledge of the specific cellular contexts in which they act. *In silico* deconvolution of bulk RNA-seq data through use of single-cell RNA-seq reference expression matrices provides a promising approach to resolve cell-type/state specific gene expression and regulatory effects, thereby advancing our understanding of the cellular mechanisms of complex traits and diseases in humans^14,16–18^. Yet, there are no such studies that have been conducted in cattle and any other livestock species.

In this study, through integrating 8,866 bulk RNA-seq samples with a bovine single-cell reference atlas^9,19^, encompassing 81 cell types derived from 22 primary tissues, we established a comprehensive atlas of cell-type specific regulatory variants in cattle, herein referred to as the CattleCell-GTEx (**Fig.1; Supplementary Table 1 and 2**). We first determined the ‘best practice’ for *in silico* bulk tissue deconvolution thorough the benchmarking of seven different algorithms in simulated datasets. For each bulk tissue, we then inferred the cell-type and cell-state compositions, and the cell-type specific transcriptional profile present, as well as systematically mapped cell-type-specific regulatory variants through a comprehensive series of approaches including: cell-type stratified eQTL (cseQTL) mapping, cell-type interaction eQTL (cell-type ieQTL) mapping, and cell-state interaction eQTL (cell-state ieQTL) mapping. Furthermore, leveraging this resource, we prioritized potential causal variants, genes, cell types, and tissues for 44 economically important production traits in cattle, and demonstrated that tissue and cell-type specific regulatory variants are significantly enriched in divergent selection signatures between dairy (*n* = 651) and beef (*n* = 552) cattle. In summary, we provided a valuable open-access resource for the cattle genetics and genomics community (https://cattlecellgtex.farmgtex.org/), and offered novel insights into the cellular molecular mechanisms underlying complex traits and selective breeding in cattle.

**Fig. 1.**
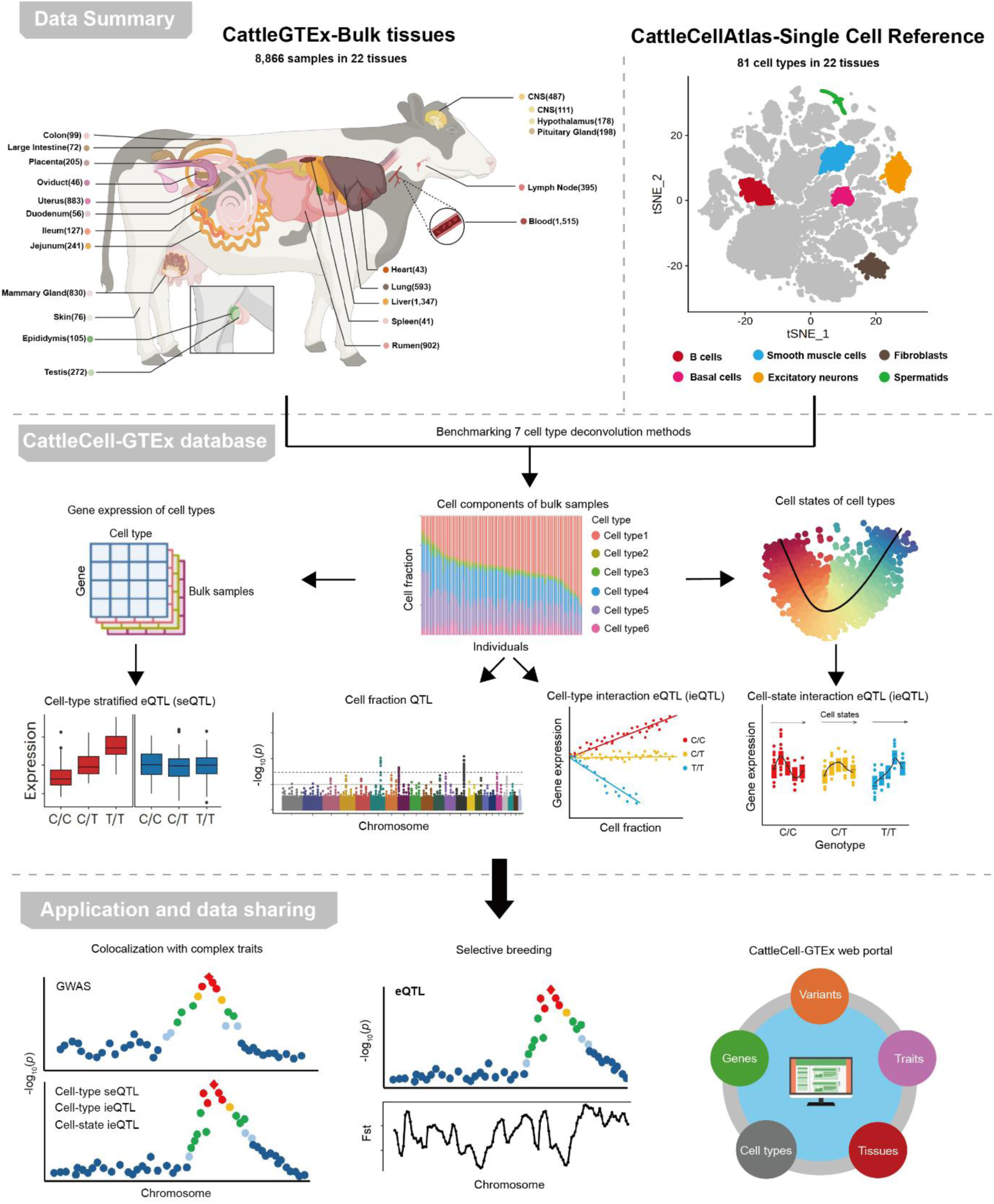
Overview of the CattleCell-GTEx project. Data summary: A total of 8,866 bulk RNA-seq samples in 22 tissues were collected from the CattleGTEx project (v1). A total of 999,192 single cells of 22 matching tissues, representing 81 distinct cell types, were collected from the Cattle Cell Atlas project. CattleCell-GTEx database: We first detected the ‘best practice’ for cell-type fraction deconvolution thorough benchmarking seven distinct algorithms in simulated datasets. We then inferred cell-type fraction, cell-state fraction and cell-type specific gene expression in all the bulk tissues, and systematically mapped regulatory variants via different approaches, including cell-type fraction QTL (fQTL), cell-type stratified eQTL (cseQTL), cell-type interaction eQTL (cell-type ieQTL), and cell-state interaction eQTL (cell-state ieQTL). Application and data sharing: Furthermore, this resource was utilized to prioritize potential causal variants, genes, cell types, and tissues for 44 complex traits of economic importance, and selection signals between dairy (n = 651) and beef (n = 552) cattle in cattle. The CattleCell-GTEx website was constructed to freely query and download all the processed data and results.

## Results

### Data summary of the CattleCell-GTEx

After removing low-quality data (**Methods**), we retained 8,866 bulk RNA-seq samples from 22 tissues with an average sample size of 506, ranging from 41 in spleen to 1,515 in blood (**Supplementary Fig. 1a**). Based on expressions of 21,385 protein-coding genes (PCGs), as expected, these bulk RNA-seq samples were clustered well with respect to their designated tissue types, indicating that the bulk RNA-seq data can reflect tissue-specific biology (**Supplementary Fig. 1b**). After genotype imputation with a multi-breed reference panel comprising of 2,742 animals from 13 breeds worldwide (**Methods**), we obtained a total of 4,317,531 high-quality common (INFO > 0.75 and MAF > 0.05) single-nucleotide polymorphisms (SNPs) for the subsequent *cis*-expression quantitative trait loci (*cis*-eQTL) mapping, where *cis*-eQTL were defined as ±1 Mb from the transcription start site (TSS) of each tested PCG. With regards to the single-cell RNA-seq (scRNA-seq) data, we obtained 999,192 high-quality single cells derived from 22 tissues present in the bulk samples (**Methods**), which represent 81 distinct cell types, from the Cattle Cell Atlas project^19^ (**Supplementary Fig. 1a**). Although cells primarily clustered by cell lineages rather than tissue types (**Supplementary Fig. 1c, 1d**), same cell types from different tissues also showed distinct expression profiles, such as CD4+ T cells from heart, lung, and uterus (**Supplementary Fig. 1e**). This underlined the importance of using single cell reference data from matching tissues for the downstream cell-type/state deconvolution analysis of bulk tissue samples.

### Deconvolution of bulk tissue samples

To identify the ‘best practice’ for bovine bulk tissue deconvolution, we benchmarked seven commonly used approaches under diverse simulated scenarios, representing three distinct distribution models (Uniform, Normal and Binormal) of cell composition in three different tissues, namely central nervous system (CNS), heart and ileum (**Fig. 2a, Supplementary Fig.2, Methods**). Among all seven evaluated methods, DWLS^20^ demonstrated the highest accuracy, the lowest Root Mean Square Error (RMSE), and the greatest robustness across all simulated scenarios for both common and rare cell types, followed by CIBERSORT^21^ and MuSiC^22^ (**Fig. 2b; Supplementary Fig.3**). Therefore, we employed DWLS to deconvolute all 8,866 bulk RNA-seq samples from 22 tissues using single-cell references of matching tissues. This approach successfully estimated cell-type fractions for 81 cell types across 22 tissues, revealing stark inter- and intra-tissue cell type composition differences (**Supplementary Fig.4-5, Supplementary Fig.6a**). For instance, macrophages had an average fraction of 10%, ranging from 0% to 81%, across 831 mammary gland samples (**Supplementary Fig.5**). In total, we observed 100 common (the estimated median fraction of a cell type ≥ 5% in a tissue) and 197 rare (< 5%) tissue-cell pairs across all 22 tissues (**Supplementary Table 3**). The rare cell types exhibited higher inter-individual variation than their common counterparts in the respective tissues (**Supplementary Fig.6b**). Furthermore, we performed trajectory analysis and then deconvoluted cell-state components (bin1-10, representing 10 continuous stages from undifferentiated to mature states) of 8,403 bulk samples for 35 cell types in 16 tissues (**Supplementary Fig.7-11; Methods**). For instance, the late state of the differentiation trajectory (bin10) in fibroblasts had an average fraction of 28%, ranging from 7% to 36%, across 198 pituitary gland samples (**Supplementary Fig.9**).

**Fig. 2.**
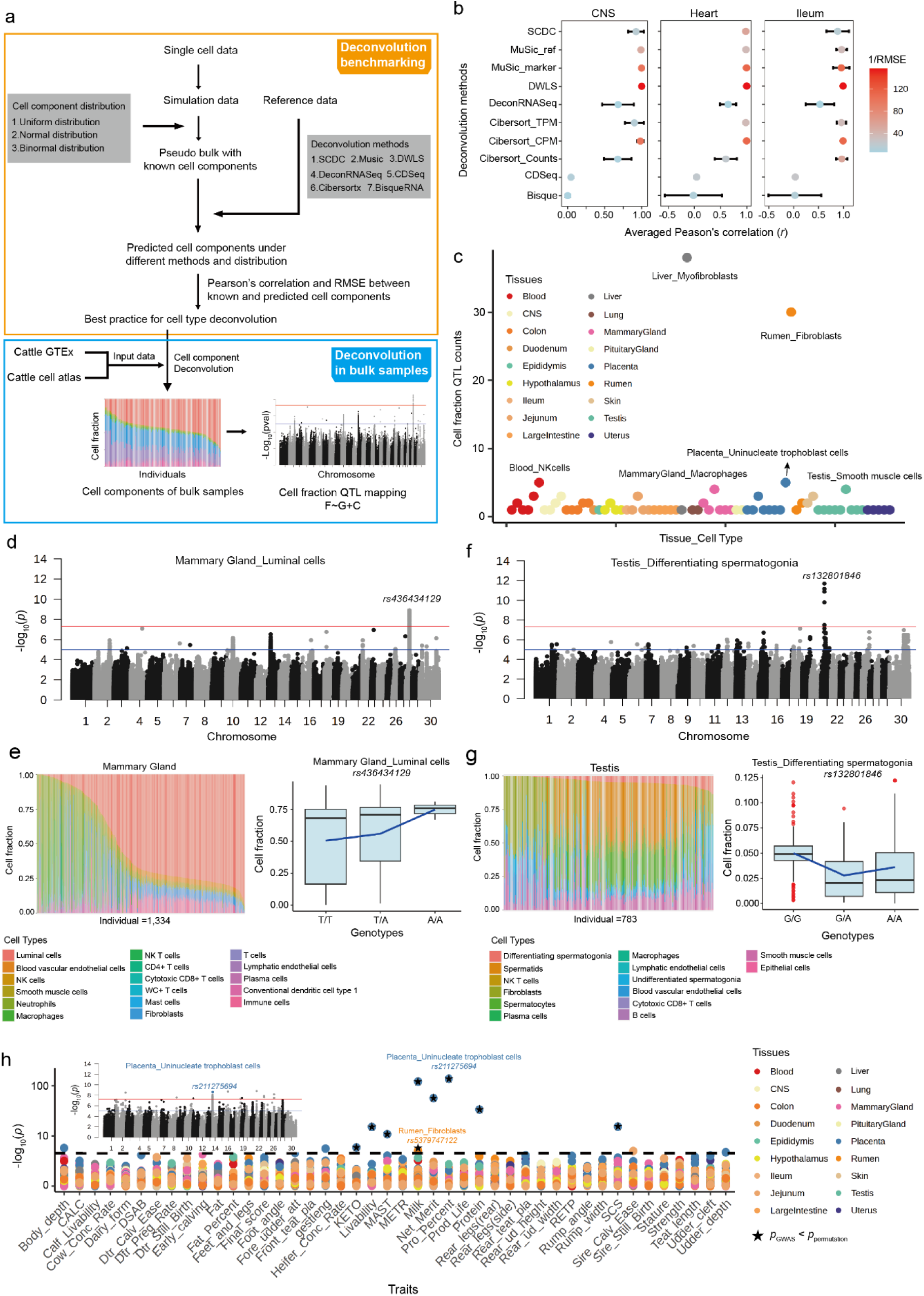
Deconvolution and cell fraction QTL (fQTL) identification across 81 cell types in 22 bovine tissues. (a) Overall workflow for cell type deconvolution and fQTL identification, including deconvolution benchmarking and deconvolution of all bulk samples. Single cell data from Cattle Cell Atlas were separated into simulated data and reference data. Pseudo bulk samples were constructed based on simulated data under three cell component distribution models. Seven deconvolution methods were then benchmarked using simulated pseudo bulk samples and single cell reference data. With the best practice, cell components of 8,866 real bulk samples were estimated and used for performing fQTL mapping between cell-type fraction and genotypes across 297 estimated tissue-cell pairs in 22 tissues. (b) Benchmarking results for seven deconvolution methods (y axis) in three tissues under multiple scenarios. The x axis represents the averaged Peason’s correlation (*r*) between true cell-type fraction and estimated fraction across simulated scenarios, while the error bar is for standard deviation (sd). The dot color represents 1/Root Mean Square Error (RMSE) value for each method. (c) Counts of fQTL across 70 tissue-cell pairs in 18 tissues. Each dot represents a tissue-cell pair, and the dot color represents tissue type. (d) Manhaton plot represents the example of fQTL in one common cell type (luminal cells of mammary gland). The red line means genome-wide significance threshold (5×10^-8^), while the blue line represents suggestive significance threshold (1×10^-5^). (e) The bar plot (left) represents cell component distribution of 17 cell types across 1,334 mammary gland samples, and the boxplot (right) represents cell fractions of luminal cells across mammary gland samples under three genotypes (T/T, T/A, A/A) of lead fQTL *rs436434129*. (f) Manhaton plot represents the example of fQTL in one rare cell types (differentiating spermatogonia of testis). The red line means genome-wide significance threshold (5×10^-8^) and the blue line represents suggestive significance threshold (1×10^-5^). (g) The barplot (left) represents cell component distribution of 14 cell types in 783 testis samples, and the boxplot (right) represents cell type fractions of differentiating spematogonias across testis samples under three genotypes (G/G, G/A, A/A) of lead fQTL *rs132801846*. The red dots represent outliners. (h) The dot plot represents the association of fQTL in 70 tissue-cell pairs with 44 cattle traits. Manhaton plot shows one fQTL example of uninucleate trophoblasts cells in placenta, which was associated with multiple traits. The black line represents the significant level with *P* = 3×10^-4^ (0.05/168). The asterisk denotes SNP–trait associations where the observed p-value was smaller than the permutation-based threshold (*p*GWAS < *p*permutation).

### The genetic control of cell-type fraction and their implications in complex traits

To investigate whether the estimated inter-individual changes in cell-type compositions are regulated by genetic variants, we conducted genome-wide association studies (GWAS) to identify cell type fraction QTL (fQTL) for the estimated 81 cell types in all 22 tissues. A total of 168 distinct fQTL (*P* < 5×10^-8^) were identified for 39 cell types in 18 tissues, representing 70 tissue-cell pairs (**Fig. 2c; Supplementary Table 4**). For instance, the estimated fraction of luminal cells, the most predominant cell type in the mammary gland, was significantly associated with a fQTL on chromosome 28 with *rs436434129* as the lead variant (**Fig. 2d, e)**. The differentiating spermatogonia, a rare cell type in the testis, was significantly associated with a fQTL on chromosome 21 with *rs132801846* as the lead variant (**Fig. 2f, g)**. There were 18 tissue-cell pairs with multiple fQTL being detected, with the most extreme cases of myofibroblasts in liver (39 fQTL) and fibroblasts in rumen (30 fQTL) (**Fig. 2c)**. By examining GWAS summary statistics of 44 complex traits, we found that *rs211275694*, the lead variant of a fQTL in uninucleate trophoblast cells of placenta, was also significantly associated with eight complex traits in cattle, including six milk production traits (e.g., milk yield, *P* = 2.1×10^-127^), cow livability (*P* = 7.5× 10^-15^) and SCS (*P* = 4.1× 10^-15^) (**Fig. 2h; Supplementary Table 5**). Additionally, the lead fSNP (*rs5379747122*) of fibroblasts in the rumen was also associated with milk yield (*p* = 8.2×10^-5^) (**Fig. 2h; Supplementary Table 5).**

### Gene expression and regulation across tissues and cell types

To explore and compare the genetic control of tissue and cellular gene expression, we first conducted *cis*-eQTL mapping at the bulk tissue level and identified that 86% of all 21,385 PCGs tested had at least one significant eQTL in at least one of 22 tissues, referred to as beGenes. The average number of detected beGenes was 5,971, ranging from 916 (5%) in skin to 11,009 (73%) in blood, which was significantly correlated with tissue sample size (**Supplementary Fig.12a; Supplementary Table 6**). Based on the similarity of eQTL effect, tissues with similar physiological functions (e.g., tissues in the digestive system) were clustered together, reflecting the tissue-specificity for gene regulation (**Supplementary Fig.12b, 12c**). To further explore cell-specific regulatory effects, we first predicted gene expressions of 49 major cell types (the estimated cell fraction > 5%) in 22 tissues using bMIND^23^, yielding 100 tissue-cell pairs (**Methods**). On average, the predicted gene expressions were significantly and positively correlated (Pearson’s *r* = 0.73) with the observed ones in the matching cell types (**Supplementary Fig.13a**). In addition, based on the predicted gene expression, most samples were clustered primarily by cell lineage rather than by tissue type, particularly within immune and epithelial cell types (**Fig. 3b; Supplementary Fig.13b-c**), which was consistent with observations from real scRNA-seq data (**Supplementary Fig. 1-1c**). Genes with cell-type-specific expression were significantly enriched in cell relevant biological processes. For instance, genes with specific expression in mammary gland macrophages were significantly enriched for immune response pathways such as IL-17 signaling pathway (FDR = 2.8×10^-6^) and cytokine-cytokine receptor interaction (FDR = 7.1×10^-6^), luminal cells for calcium signaling (FDR = 1.7×10^-4^), and fibroblasts for the extracellular matrix pathway (FDR = 6.2×10^-3^). The average *cis*-heritability of PCGs was 0.25 across 49 cell types in 22 tissues, which was significantly higher than the *cis*-heritability of genes estimated in bulk tissues (0.17) (**Supplementary Fig.13d**). Collectively, these results indicated that the predicted gene expression reflected cell-specific biological processes, and the presence of inter-individual variation in mRNA abundance was sufficient to enable subsequent cell-type-specific *cis*-eQTL mapping.

**Fig. 3.**
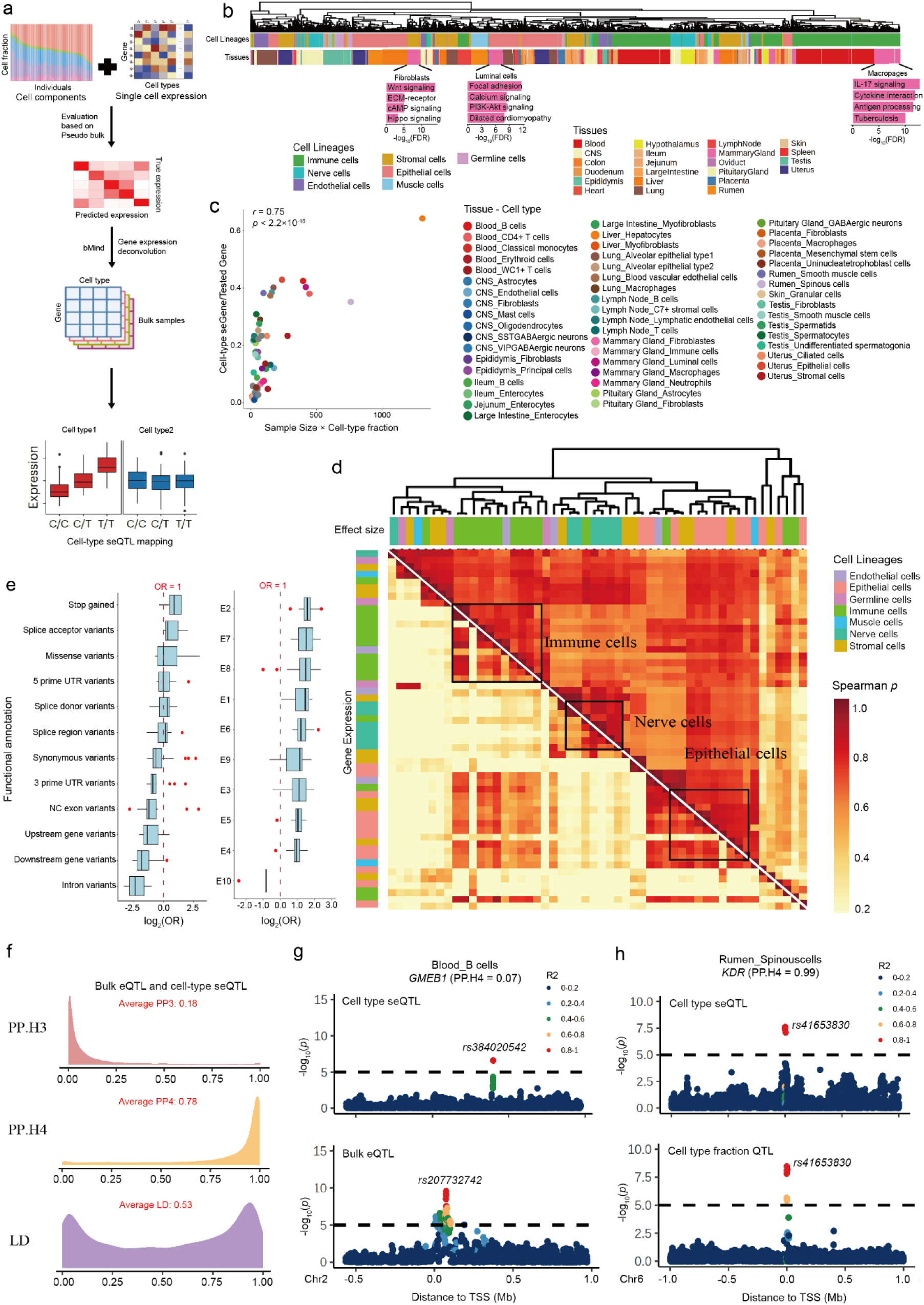
Cell type stratified eQTL (cseQTL) mapping across 100 common tissue-cell pairs. (a) Overall workflow for cseQTL mapping. Gene expressions in 100 common tissue-cell pairs were estimated based on deconvoluted cell components and gene expression from single cell reference data, which were then used for cell-type stratified eQTL (cseQTL) mapping. (b) Clustering of 35,030 tissue-cell-sample trios based on 898 differential expressed genes (adjust.p < 0.05 and log2FC > 0.5) across 100 tissue-cell pair. Samples were annotated by cell lineages (top) and tissue types (bottom). (c) Pearson’s correlation (*r*) between the proportion of detectable cseGenes and the product of sample size and cell-type fraction across 53 tissue-cell pairs with cseGenes being detected. The two-sided *P* values were obtained by Pearson’s correlation (*r*) test. (d) Heatmap of tissue-cell pairs depicting the corresponding pairwise Spearman’s correlation (*p*) of gene expression (down) and cseQTL effect sizes (up). Tissue-cell pairs are grouped by hierarchical clustering. (e) Enrichment of lead cseQTL across sequence ontology (i.e., variant types annotated by SnpEff software) and 10 chromatin states (E1: Active Transcription Start Site (TSS), E2: Flanking Active TSS, E3: Weak TSS, E4: Active Enhancer, E5: Weak Enhancer, E6: Poised Enhancer, E7: Bivalent TSS, E8: Bivalent Enhancer, E9: Repressed Polycomb, E10: Quiescent/Low Signal). (f) Colocalization between cseQTL and fQTL. The top plot represents Posterior Probability for Hypothesis 3 (PP3) value distribution with an average equal to 0.18, the middle plot represents PP4 value distribution with an average of 0.78, and the bottom plot represents Linkage Disequilibrium (LD) with an average of 0.53 between lead eSNPs in cseQTL and beQTL. (g) An example of a cseQTL for *GMEB1* identified in blood B cells. The lead variant *rs384020542* was significantly associated with *GMEB1* expression in B cells but not in bulk tissue (beQTL, lead variant *rs207732742*), with a colocalization PP4 of only 0.07. (h) An example of a cseQTL for *KDR* in rumen spinous cells, with strong colocalization with the corresponding cell-type fraction QTL, both driven by the same lead variant *rs41653830* (PP4 = 0.99).

The *cis*-eQTL mapping revealed that 85% of all 19,165 tested PCGs had at least one significant variant identified in at least one of 53 tissue-cell pairs, referred to as cell-type stratified eGenes (cseGenes) (**Fig. 3a)**. The average number of detected cseGenes across tissue-cell pairs was 2,074, ranging from 9 in enterocytes of large intestine to 7,578 in hepatocytes of liver (**Supplementary Fig.14a; Supplementary Table 7**). The proportion of cseGenes identified was positively correlated with the product of tissue sample size and cell-type fraction (**Fig. 3c**). Compared to beGenes, we identified an average of 816 novel genes as cseGenes across tissues, ranging from 4 in skin to 1,737 in testis, with the count positively correlated with sample size (**Supplementary Fig.14b**). The effect sizes of lead variants of novel cseGenes were significantly smaller than those of beGenes, indicating that a higher statistical power can be observed from mapping eQTL at the cellular level compared to the bulk tissue level, despite leveraging predicted gene expression levels (**Supplementary Fig.14c**). The conditional and fine-mapping analyses revealed that 86% and 67% of cseGenes had at least one independent cseQTL and credible set, respectively, and among these 942 were novel compared to beQTL (**Supplementary Fig. 14e-g**). Cell types were clustered primarily by cell lineages rather than tissue types according to the similarity of effect sizes of lead cseQTL (**Fig. 3d**). This result suggested that cseQTL captured intrinsic cellular regulatory mechanisms. Compared to beQTL, cseQTL were more likely to be cell-type-specific (**Supplementary Fig.8b; Supplementary Fig.14d**). A total of 74% of 8,061 cseGenes were colocalized with their corresponding beGenes (Posterior Probabilities Hypothesis 4-PP4 > 0.8), the lead variants of both with an average linkage disequilibrium (LD, r^2^) of 0.53 (**Fig. 3f**). However, 1,903 (11%) of cseGenes displayed distinct signals (PP4 < 0.25 and r^2^ < 0.2) compared to beGenes (**Fig. 3f**). For instance, in B cells of blood, the lead cseQTL of *GMEB1* was distinct from the lead beQTL (PP4 = 0.07), with a low LD (r^2^ = 0.003) (**Fig. 3g**). Overall, these findings suggested that mapping eQTL using predicted gene expression at cell type resolution can reveal novel regulatory variants compared to bulk tissue eQTL. Additionally, 21 out of 168 fQTL were colocalized with cseQTL (PP4 > 0.8; **Supplementary Table 8**). For example, an fQTL identified for spinous cells deconvolved from rumen tissue were colocalized (PP4 = 0.99) with a cseQTL of the *KDR* gene, which encodes vascular endothelial growth factor and induces endothelial proliferation, survival, migration, tubular morphogenesis and sprouting^24^, with the lead SNP, *rs41653830*, located on chromosome 6 (**Fig. 3h**).

Similar to beQTL, lead cseQTL were highly enriched in stop-gained variants, missense variants, 5’ UTRs, and splice donor/acceptor sites^9,11,12^ (**Fig. 3e**, **Supplementary Fig.15a**). Among 10 chromatin states predicted previously in cattle^25^, the lead cseQTL showed a higher enrichment in promoters than enhancers, with the highest enrichment in flanking TSS regions (**Fig. 3e; Supplementary Fig.15b-c**). By examining single-cell ATAC-seq data from six mammary gland samples, we assessed the enrichment of lead cseQTL within 14,876 cell-type differential open chromatin regions across five cell types, including fibroblasts, luminal hormone-responsive (HR) cells, luminal secretory (Sec) cells, macrophages and T cells (**Supplementary Fig.15d; Supplementary Table 9**). The cseQTL exhibited the strongest enrichment in the open chromatin regions specific to their corresponding cell types, particularly for luminal cells and fibroblasts, providing additional evidence that cseQTL can capture cell-type-specific gene regulation mechanisms (**Supplementary Fig.15e**).

### Cell-type/state interaction *cis*-eQTL mapping

To further explore cell-specific regulatory effects, we conducted a complementary analysis to detect the interaction effects of genotype × cell type fraction on gene expression, referred to as cell-type interaction eGenes (ieGenes). Analogous to cseGenes, the discovery power of cell-type ieGenes was positively correlated with the product of sample size × cell-type fraction (**Fig. 4a**). In total, we detected 16,341 cell-type ieGenes, representing 76% of all tested PCGs, across 46 tissue-cell pairs, ranging from 62 of NK T cells in jejunum to 7,032 of CD4+ T cells in blood (**Supplementary Fig.16; Supplementary Table 7**). For instance, the regulatory effect of *rs4968146* on the expression of *FLOT1* was significantly interacted with the enrichment of CD4+ T cell in blood (**Fig. 4b**). Compared to beQTL, 76% of cell-type ieGenes appeared to be driven by distinct genetic variants (PP4 < 0.25 and LD < 0.2), suggesting that cell-type ieQTL mapping can capture additional regulatory signals that are masked in bulk tissue. Moreover, cell-type ieGenes showed a significantly higher enrichment in tissue-specific beGenes than shared ones (*P* = 7.4×10^-7^; **Fig. 4c**). For instance, among 6,957 ieGenes detected in fibroblasts from 10 different tissues, 10,009 (88%) of 11,320 lead cell-type ieQTL were characterized in only one of these tissues, whereas 3,150 (13%) of 24,116 lead beQTL were in the same set of tissues (**Fig. 4d; Supplementary Fig.17a)**. On average, 98% of cell-type ieGenes were also identified as cseGenes across 46 tissue–cell pairs. However, the colocalization analysis revealed that 67% of cell-type ieGenes appeared to be driven by distinct genetic variants compared to cseGenes (PP4 < 0.25 and LD < 0.2; **Fig. 4e**). These findings suggested that cell-type ieQTL mapping can capture additional regulatory signals that were missed by cseQTL mapping. Notably, we observed a stronger correlation in effect sizes between cell-type-specific ieQTL and their matched cseQTL, compared to non-matching cell types, especially in tissues with large sample sizes such as blood and mammary gland (**Fig. 4f-g; Supplementary Fig.17b**). For example, the lead ieQTL (*rs519268026*) of *ENSBTAG00000050728* in CD4+ T cells of blood was also identified as the lead cseQTL in CD4+ T cells but was not significant in any other cell types like B cells (**Fig. 4h**). Furthermore, compared to all cell-type ieQTL, the cell-type-specific ieQTL were also more likely to be colocalized with cseQTL (**Supplementary Fig.17c**), providing additional evidence that both cell-type ieQTL and cseQTL mapping can capture cell-type-specific regulatory effects.

**Fig. 4.**
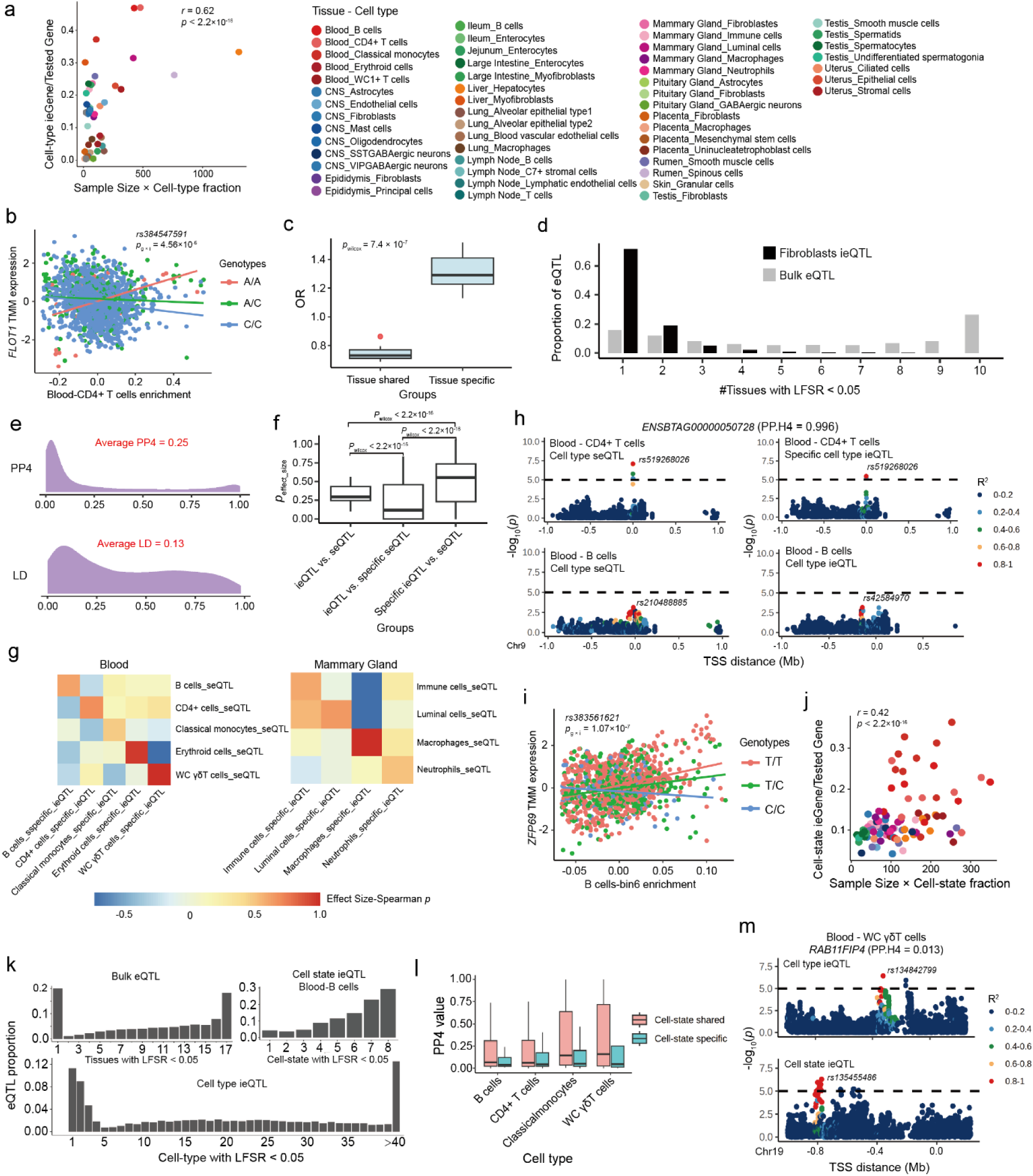
Cell-type/cell-state interaction eQTL (ieQTL) mapping across 100 common tissue-cell pairs. (a) Pearson’s correlation (*r*) between the proportion of detectable cell-type ieGenes and the product of sample size and cell-type fraction across 44 tissue-cell type pairs with cell-type ieGenes being detected. The two-sided *P* values was obtained by Pearson’s correlation (*r*) test. (b) Effect of cell-type ieQTL (*rs384547591*) of *FLOT1* interacted with CD4+ T cells enrichment in blood. The two-sided *P* value was calculated by the linear mixed regression model. The lines were fitted by a linear regression model using the geom_smooth function from ggplot2 (v3.3.2) in R (v4.0.2). The color of lines represents three genotypes of *rs384547591* (A/A, A/C, C/C). (c) Enrichment level (odd ratio) between cell-type ieQTL and tissue-specific/shared bulk eQTL (beQTL) identified by MashR (v0.2.6). The two-sided *P* value was calculated by the Wilcox test. (d) The overall tissue-sharing pattern of beQTL and fibroblasts ieQTL at Local False Sign Rate (LFSR) < 5% across 10 tissues. (e) Colocalization results between cell-type stratified eQTL (cseQTL) and cell-type ieQTL. The top plot represents the PP4 distribution and the bottom one represents the Linkage Disequilibrium (LD) of lead SNP between cseQTL and cell-type ieQTL. (f) Pairwise spearman’s correlation (*ρ*) of effect sizes between cell-type ieQTL and cseQTL, cell-type specific ieQTL and matched cseQTL, and cell-type specific cseQTL and matched cell-type ieQTL in the same tissue-cell pairs. The two-sided *P* value was calculated by the Wilcox test. (g) Pairwise spearman’s correlation (*ρ*) of effect sizes between cseQTL and cell-type specific ieQTL across cell-types of blood (left) and mammary gland (right). (h) Manhaton plot represents an example of colocalization between cseQTL and cell-type specific ieQTL in the blood. This colocalization signal was specifically observed between B cell cseQTL and ieQTL, but was absent between B cell cseQTL and CD4+ T cell ieQTL. (i) Effect of cell-state ieQTL (*rs383561621*) of *ZFP69* interacted with cell-state bin 6 of B cells enrichment in blood. The two-sided *P* value was calculated by the linear mixed regression *cis*-ieQTL model. The lines were fitted by a linear regression model using the *geom_smooth* function from ggplot2 (v3.3.2) in R (v4.0.2). The color of lines represents three genotypes of *rs383561621* (T/T, T/C, C/C). (j) Pearson’s correlation (*r*) between the proportion of detectable cell-state ieGenes and the product of sample size and cell-state fraction across 21 tissue-cell type pairs. The two-sided *P* values were obtained by Pearson’s correlation (*r*) test. (k) The tissue/cell-type/cell-state sharing patterns of eQTL under bulk eQTL, cell-type and cell-state ieQTL levels. (l) PP4 value distribution of colocalized eGenes between cell-type ieGenes and cell-state specific or shared ieGenes across four cell-types (i.e., B cells, CD4+ T cells, Classical monocytes and WC+ γδT cells) of blood. (m) The Manhatton plot represents a novel ieQTL (*rs134842799*) in *RAB11FIP4* identified by cell-state ieQTL in WC+ γδT cells of blood, but not observed at cell-type ieQTL in the same tissue-cell pair (PP4 = 0.013).

To explore the interaction effects of genotype × cell states on gene expression, we further performed cell-state interaction eQTL (cell-state ieQTL) mapping based on the estimated cell states above, and identified 67% of 21,385 tested PCGs as cell-state ieGenes in at least one of 21 tissue-cell pairs, ranging from 6% in endothelial cells of CNS to 63% in classical monocytes of blood (**Supplementary Fig.18a; Supplementary Table 7**). For example, the regulatory effect of *rs383561621* on *ZFP69* expression was significantly interacted with B cell states in blood (**Fig. 4i**). The discovery power of cell-state ieGenes was positively correlated with the product of sample size × cell-state fraction (**Fig. 4j; Supplementary Fig.18b**). Compared to the three types of eQTL detected above, including beQTL, cseQTL, and cell-type ieQTL, although cell-state ieQTL only identified an average of two additional ieGenes across 21 tissue-cell pairs, the colocalization analyses of cell-state ieQTL with all other eQTL revealed 12,680 (35%) additional regulatory variants (PP4 < 0.25 and LD < 0.2). Analysis of the sharing patterns of cell-state ieQTL highlighted that cell-state ieQTL were more likely to be shared among cell states compared to the tissue or cell-sharing patterns of beQTL, cseQTL, and cell-type ieQTL (**Fig. 4k; Supplementary Fig.14d; Supplementary Fig.18c**). Compared to cell-state shared ieQTL, cell-state specific ieQTL were less likely to be colocalized with cell-type ieQTL, thereby enhancing the discovery of novel cell-state specific regulatory effects (**Fig. 4l**). For example, in WC+ γδT cells derived from blood, the gene *RAB11FIP4* exhibited distinct lead ieQTL at the cell-state and cell-type levels (**Fig. 4m**).

### Deciphering regulatory mechanisms of complex traits

To investigate the regulatory mechanisms underlying complex traits in cattle, we systematically integrated all four types of eQTL detected above, including 18,391 beGenes, 16,290 cseGenes, 16,341 cell-type ieGenes, and 14,328 cell-state ieGenes, with 2,070 independent loci for 44 complex traits discovered by GWAS that represented 22 body conformation, 6 milk production, 8 reproduction, and 8 health traits (**Supplementary Fig.19**). These GWAS loci exhibited significant enrichments in all types of eQTL, with the strongest enrichment observed in cseQTL, followed by cell-type/state ieQTL, and finally beQTL (**Fig. 5a**). To prioritize the potential causal variants and genes for complex traits, we colocalized each of 2,070 GWAS loci with all types of eQTL. In total, 75% (1,555) of GWAS loci, ranging from 33% (4) in calving to conception interval to 96% (97) in milk protein percentage, were colocalized (PP4 > 0.8) with at least one type of eQTL (**Fig. 5b; Supplementary Fig.20b**), whereas beQTL contributed to 48% (994) only (**Supplementary Fig.20a**). Among all 1,555 colocalized GWAS loci, 1,120 (72%) were not colocalized with the nearest genes of the lead GWAS SNP (**Fig. 5c**), indicating the complexity of gene regulation underlying complex traits. The four types of eQTL showed their unique contributions to the GWAS colocalization, where cell-state ieQTL contributed to the most, with 131 unique colocalization, followed by cell-type ieQTL (35), beQTL (35), and finally cseQTL (23) (**Fig. 5b**). This suggested the importance of mapping regulatory variants at cell-type/state level for deciphering the molecular function of genomic variants associated with complex traits in cattle.

**Fig. 5.**
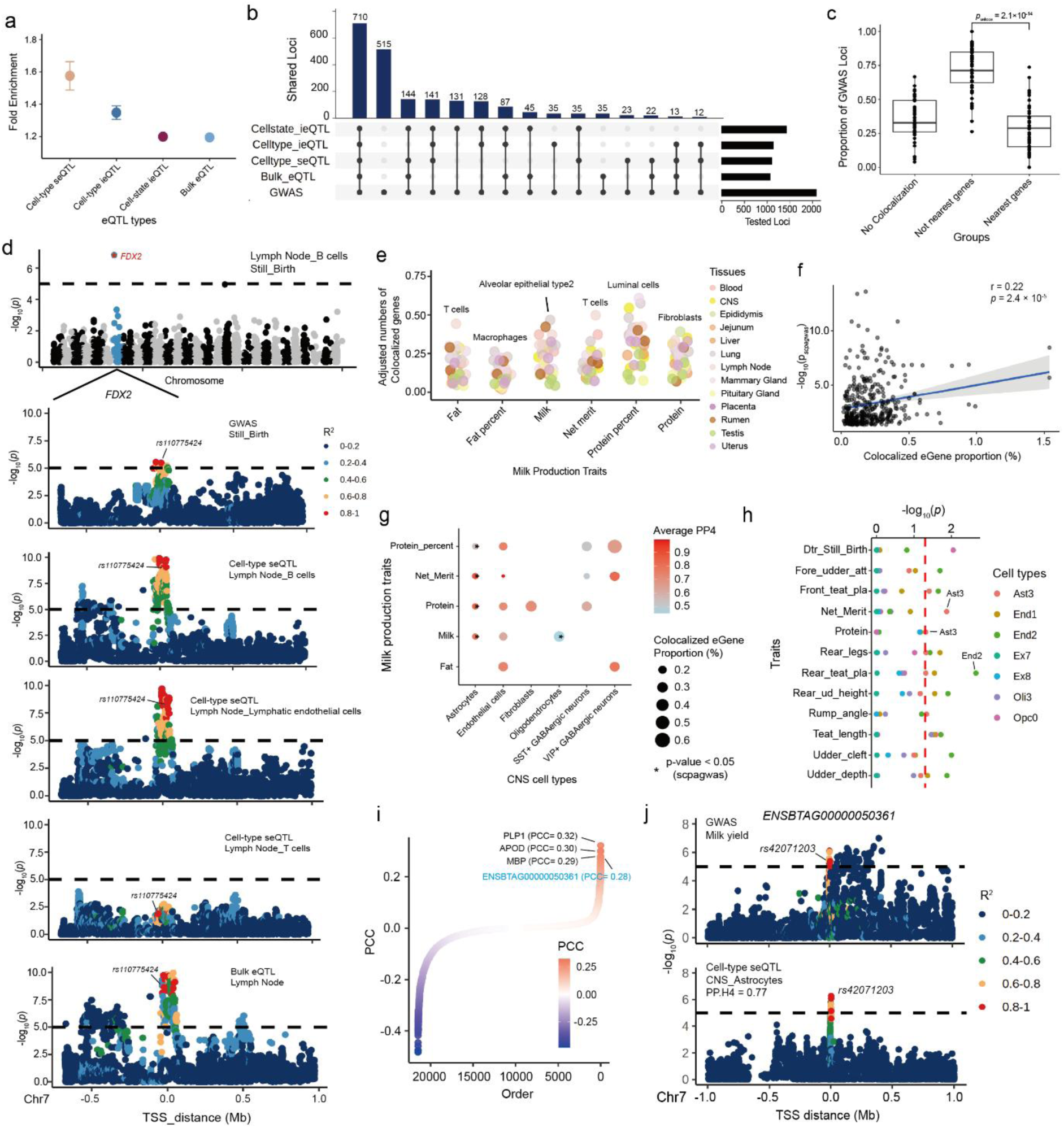
Deciphering regulatory mechanisms of 44 complex traits in cattle. (a) Enrichment between four classes of eQTL (beQTL, cseQTL, cell-type and cell-state ieQTL) and genome-wide associations (GWAS) of 44 cattle traits using QTLenrich (v.2.0). (b) The number of colocalized GWAS loci by each eQTL (beQTL, cseQTL, cell-type and cell-state ieQTL). Each blue bar represents a combination of different eQTL. Each black bar (right) represents the number of GWAS loci colocalized in the respective combination. (c) Proportion of three types of GWAS loci regarding the colocalization results, where 44 GWAS traits are shown in each category. No colocalization, GWAS loci that are not colocalized with any eGenes in 44 tissues. Not nearest gene, GWAS loci whose colocalized eGenes are not nearest genes to GWAS lead SNPs. Nearest gene, GWAS loci whose colocalized eGenes are the nearest ones. Each dot represents a complex trait. The two-sided *P* values were obtained by the Wilcox test. (d) The association of *FDX2* with stillbirth. The top Manhattan plot represents the transcriptome-wide association (TWAS) results of stillbirth in the lymph node. The four following Manhattan plots show the colocalization of stillbirth GWAS (top), cell-type seQTL in B cells (second), cell-type seQTL in lymphatic endothelial cells (third) and bulk eQTL (bottom) of *FDX2* on chromosome 7 (chr 7) in the lymph node. (e) The number of colocalized cseGene per 100 cseGenes in each tissue-cell-trait trio between six milk production traits (i.e., milk yield, fat yield, fat percent, protein yield, protein percent, and net merit) and cseQTL. (f) Pearson’s correlation (*r*) between significant levels of cell-type enrichment analysis and proportion of colocalized cseGenes in all cseGenes, across 44 traits and 53 tissue-cell pairs. The two-sided *P* values were obtained by Pearson’s correlation (*r*) test. (g) The enrichment results and colocalization between six cell-types in Central Nervous System (CNS) and five milk production traits. The dot size represents colocalized cseGene counts, and the dot color represents the average PP4 of colocalization. The asterisk represents the enriched cell-type also had colocalized cseGenes in the respective trait. (h) The significant enrichment results between seven cell-types from cattle brain spatial data and 12 complex traits. The red line represents the significant level with *p* = 0.05. (i) Pearson’s correlation (PCC) between milk yield and each of 21,385 genes in astrocytes of CNS. (j) Manhattan plots showing the –log₁₀(p) distribution of association signals for milk yield (GWAS) and cseQTL in CNS astrocytes across genetic variants within the locus of *ENSBTAG00000050361*. Variants are colored according to their linkage disequilibrium (LD, r²) with the lead SNP (*rs42071203*). The top panel displays GWAS association signals; the bottom panel shows astrocyte-specific cseQTL signals. The lead variant exhibits strong colocalization between the two datasets (PP.H4 = 0.77).

For example, *rs383773615*, a cseQTL for *ENSBTAG00000051883* in T cells of lymph node, exhibited a strong colocalization with calving ease (PP4 = 0.82) and livability (PP4 = 0.81). In contrast, no colocalization signal was observed for the beQTL of *ENSBTAG00000051883* in lymph node (**Fig. 6a**). Although it remains a functionally uncharacterized gene in the bovine genome, *ENSBTAG00000051883* shares homology with zinc finger transcription factors (ZNFs), a gene family implicated in a variety of biological processes and diseases, including human cancers, embryonic lethality, and neurological disorders^26^ (e.g., *ZNF775* in humans and *ZNF517* in chimpanzee). Notably, *ENSBTAG00000051883* is located approximately 0.1 Mb from the bovine *ZNFO* gene, which has been linked to embryonic development in cattle^27^. In another example involving cell-type ieQTL, the lead variant *rs136273564* of *KLHL18*, showed a strong colocalization with somatic cell score (SCS) (PP4 = 0.82) specifically in hepatocytes of the liver. This gene was previously shown to be involved in inflammation regulation via ubiquitin-mediated immune signaling^28^. However, this colocalization signal was not detected in the corresponding cseQTL (PP4 = 2.2×10^-5^) or beQTL (PP4 = 2.6×10^-5^) (**Fig. 6b**). Additionally, a cell-state ieQTL—*rs111015829* in *GLTPD2*, was colocalized with the calving ease trait (PP4 = 0.95) in hepatocytes. This gene plays a role in glycolipid-mediated membrane signaling and immune cell regulation^29^. This colocalization signal was absent in the corresponding cseQTL and cell-type ieQTL (**Fig. 6c**).

**Fig. 6.**
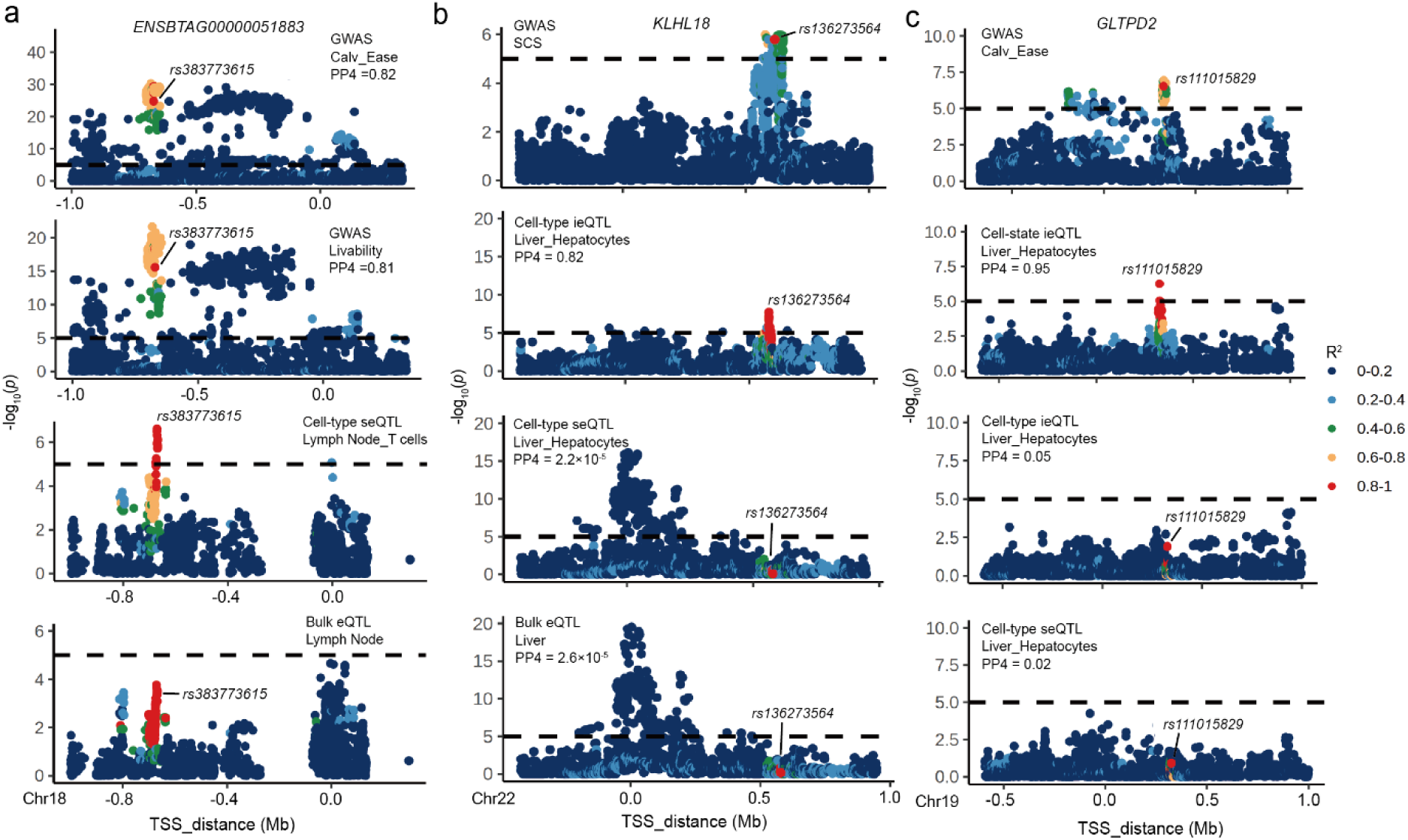
Three examples of specific colocalization between three types of cell-resolved eQTL and GWAS loci. (a) The Manhattan plot represents a cseQTL for *ENSBTAG00000051883* in T cells of lymph node, exhibited a strong colocalization with calving ease (PP4 = 0.82) and livability (PP4 = 0.81). In contrast, no colocalization (PP4 = 0.02) was observed for the beQTL of *ENSBTAG00000051883* in lymph node. (b) The Manhattan plot represents one cell-type ieQTL, the lead variant *rs136273564* of *KLHL18*, showed a strong colocalization with somatic cell score (SCS) (PP4 = 0.82) specifically in hepatocytes of the liver. However, this colocalization was not detected in the corresponding cseQTL or beQTL. (c) The Manhattan plot represents a cell-state ieQTL—*rs111015829* in *GLTPD2*, was colocalized with calving ease trait (PP4 = 0.95) in hepatocytes, but was absent in the corresponding cseQTL and cell-type ieQTL.

Furthermore, we conducted cell-type transcriptome-wide association studies (TWAS) for all 44 complex traits (**Supplementary Fig.21a; Methods**) and detected 3,823 significant gene-cell-trait trios (FDR < 0.05, **Supplementary Fig.21b**). For example, *FDX2* was the top associated gene in the TWAS analysis of stillbirth, which was identified in B cells of the lymph node. The *FDX2* gene has been shown to be involved in various nutritional metabolic processes^30^ and the cseQTL (*rs110775424*) associated with this gene in lymph node B cells was also colocalized with stillbirth (PP4 = 0.74). This colocalization was exclusively observed in B cells and lymphatic endothelial cells of lymph node, but not in T cells derived from the same tissue—likely because *FDX2* was not identified as a cseGene in T cells (**Fig. 5d**). Unlike T-cells, which primarily mediate cellular immunity, B-cells are centrally involved in antibody production and immune tolerance—processes that are integral to pregnancy^31,32^. These results indicated a potential mechanistic role of humoral immunity in phenotypic variation for stillbirth.

To detect trait-relevant tissues and cell types, we first calculated the number of colocalized cseGenes per 100 cseGenes across 53 tissue-cell pairs. Immune cells contributed to the most GWAS colocalization among all 44 complex traits (**Supplementary Fig.20c**). Luminal cells in the mammary gland exhibited the most colocalization signals from milk protein percentage **(Fig. 5e)**, consistent with the function of luminal cells in nutrient transport and milk secretion^33^. Within the lymph node, compared to other cell types, immune cells (e.g., T cells and macrophages) also showed stronger colocalizations with several milk production traits like fat yield and net merit **(Fig. 5e)**. This was consistent with that immune responses can influence the integrity of the blood-milk barrier (BMB), thereby modulating milk composition and secretion^34^. Furthermore, we conducted a complementary analysis to detect trait-relevant cell types through testing whether GWAS signals of a trait are significantly enriched in genes with cell-type specific expression across all 250 tissue-cell pairs, which were obtained from the Cattle Cell Atlas dataset^19^. In total, we identified 2,586 significant (FDR < 0.05) cell-tissue-trait trios between 44 traits and 244 tissue-cell pairs, ranging from 8 tissue-cell pairs in fat percentage to 60 in metritis resistance (**Supplementary Fig.22a**). The significance of the trait-cell-tissue relationships detected by the enrichment analyses were positively correlated (Pearson’s *r* = 0.22, *P* = 2.4 × 10^-5^) with the proportion of colocalized cseGenes (**Fig. 5f**). More specifically, 214 significant cell-tissue-trait trios detected by enrichment analysis were also supported by GWAS-cesQTL colocalization analysis (**Supplementary Fig.22b**). Astrocytes in the CNS demonstrated a higher enrichment degree and more colocalized genes for four milk production traits (protein percentage, net merit, protein yield, and milk yield), compared to other traits. This was consistent with a previous report that neuroregulatory mechanisms played a prominent role in lactation through neuronal-glial dialogue^35,36^ (**Fig. 5g; Supplementary Fig.22b**). To further investigate the spatial association of brain cell types with milk production, we constructed a spatial cell atlas of the cattle brain, revealing seven distinct cell types in three major brain regions (i.e., neocortex, pallidum and white matter) (**Supplementary Fig.23**). Among these, astrocytes, located at white matter region in the brain, were significantly associated with milk production traits such as net merit and protein yield (**Fig. 5g-h**). Notably, the cseQTL of *ENSBTAG00000050361*, which has a positive Pearson’s correlation coefficient of 0.28 with milk yield based on enrichment analysis in astrocytes (**Fig. 5i; Supplementary Table 10**), was also colocalized with milk yield (PP4 = 0.77; **Fig. 5j**). Although it is a novel gene in the cattle genome, *ENSBTAG00000050361* is orthologous to *APOBR* in other mammals, which has previously shown to be related to lipids, lipid-soluble vitamin, and other nutrients metabolic processes^37^, further supporting the potential importance of astrocytes or related cell types in lactation and milk production.

### Understanding regulatory mechanisms underlying adaptive selection and evolution

To elucidate the regulatory mechanisms underlying selective breeding, we scanned genome-wide selection sweeps between 651 Holstein dairy cattle and 552 beef cattle via an *F*_ST_-based approach (**Supplementary Table 11**). The total of 248,690 genomic regions tested were evenly divided into 10 deciles (*F*_ST_1-10) based on their *F*_ST_ values from the smallest (no selection) to the largest (strongest selection) (**Supplementary Fig.24a**). For example, genomic regions overlapped with *DDX56* (Fst = 0.6), *CCND2* (*F*_ST_ = 0.53), *RPL13* (*F*_ST_ = 0.51) and *SPG7* (*F*_ST_ = 0.5) were under the strongest selection group (*F*_ST_10) between dairy and beef cattle and were also reported in previous studies, such as *DDX56* was selected between three South African cattle breeds and participated in immune response in cattle, whereas *CCND2* was selected among five Swedish cattle breeds and affected cattle body size and stature^38–41^ (**Fig. 7a**). The enrichment of the fine-mapped beQTL in each of 10 deciles was then tested (**Supplementary Fig.24b, 24c**). In general, the tissue-shared beQTL was more likely to be enriched in both *F*_ST_1 and *F*_ST_10 compared to the rest of the genome (**Fig. 7b**). Compared to tissue-shared beQTL, tissue-specific beQTL showed a significantly higher enrichment in regions with larger *F*_ST_ values, particularly in *F*_ST_10 (**Fig. 7b**). Among all the examined 17 tissues, tissue-specific beQTL of large intestine, pituitary gland, hypothalamus, and jejunum, displayed the highest enrichment in the *F*_ST_10 group, suggesting that the digestive and nervous systems have played important roles in the divergent breeding of dairy and beef cattle (**Fig. 7c; Supplementary Fig.24d**). We further considered 3,198 beGenes residing in *F*_ST_10 as potentially selected genes between dairy and beef cattle, ranging from 42 in the large intestine to 759 in the liver (**Fig. 7c**). The selected beGenes in blood and liver were significantly enriched in pathways related to milk production, such as the PI3K-Akt signaling pathway (FDR = 6×10⁻⁴) and steroid hormone biosynthesis (FDR = 8×10⁻⁵), suggesting their potential roles in the genetic improvement of milk production during selective breeding of dairy cattle (**Fig. 7c; Supplementary Fig.24e**). This was further supported by that GWAS loci of milk production showed a higher enrichment in genomic regions with higher *F*_ST_ (**Fig. 7d**). Compared to other tissues, blood and liver had more colocalized beGenes (3 in blood and 10 in liver; PP4 > 0.8) with milk production (**Fig. 7e**). For example, the lead beQTL (*rs211403999*) of *FBXL6*, a causal gene for mastitis and milk production traits in the liver^42^, was colocalized with Net Merit (PP4 = 0.95) and resided in a strong selection region (*F*_ST_ = 0.25; **Fig. 7f**). The frequency of C allele of *rs211403999* was elevated in the Holstein dairy cattle compared to beef cattle (**Fig. 7g**). Another example was *AKR1B1*, whose lead beQTL (*rs137649450*) in liver was colocalized with milk protein percentage (PP4 = 0.93) and located in a region of high selection (*F*_ST_ = 0.17; **Fig. 7h**). This gene family plays pivotal roles in the biosynthesis and metabolism of steroid, and influences the production of steroid hormones such as androgens, estrogens, and progesterone, which are critical regulators of lactation^43^.

**Fig. 7.**
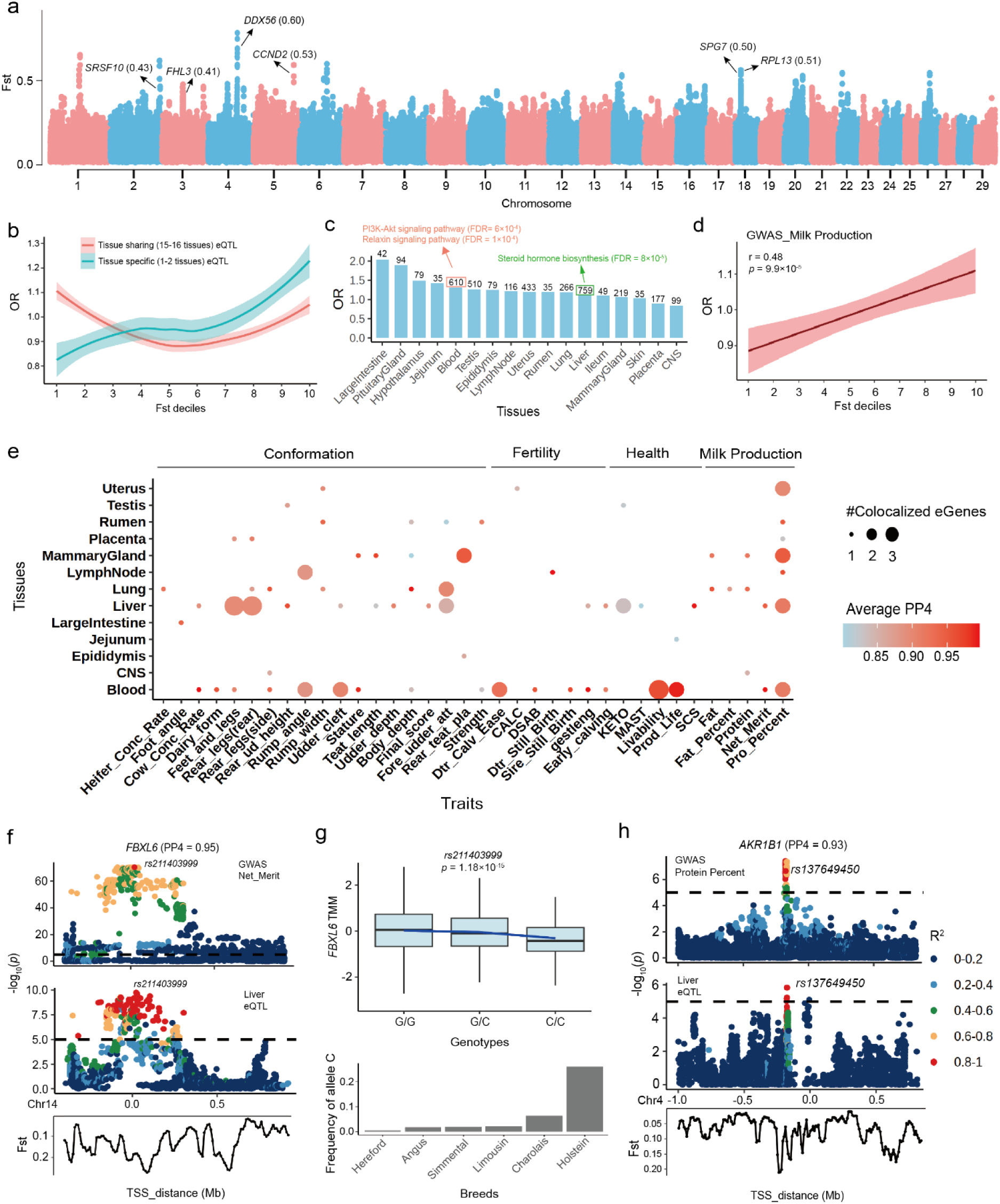
Liking bulk eQTL to selection signatures between dairy and beef cattle. (a) Genome-wide *F*_ST_ distribution across 248,690 genomic regions. Genes overlapping with the top 10% *F*_ST_ regions (putative targets of strong selection) are highlighted, including previously reported loci such as *SRSF10*, *FHL3*, *CCND2*, *SPG7*, *RPL13*, and *DDX56*. (b) Enrichment (odds ratio) of shared and tissue-specific beQTL across 10 *F*_ST_ deciles. The lines were fitted by using the geom_smooth function from ggplot2 (v3.3.2) in R (v4.0.2). (c) The number of tissue-specific beGenes under strong selection (top 10% *F*_ST_) across 17 tissues. Associated KEGG pathways related to milk production are indicated, including the PI3K-Akt signaling (FDR = 6×10^-4^), relaxin signaling (FDR = 1×10^-4^), and steroid hormone biosynthesis pathways (FDR = 8×10^-5^). (d) Enrichment of GWAS loci for five milk production traits across the 10 *F*_ST_ deciles. The lines are fitted by a linear regression model using the geom_smooth function from ggplot2 (v3.3.2) in R (v4.0.2). The two-sided *P* values are obtained by Pearson’s correlation (*r*) test. (e) The bubble plot comparing the number of beGenes that were both colocalized with complex traits and under strong selection. Dot size reflects the number of both selected and colocalized genes per trait, while dot color indicates the average colocalization posterior probability 4 (PP4). (f) Manhattan plots illustrating an example of a both selected and colocalized beGene (*FBXL6*) in liver, associated with net merit. The beQTL *rs211403999* shows high LD with the trait signal and lies in a high-*F*_ST_ region (*F*_ST_ = 0.25). (g) Functional interpretation of *rs211403999*: boxplot (top) shows expression of *FBXL6* across three genotypes (G/G, G/C and C/C), while the barplot (bottom) shows the frequency distribution of the allele C across five cattle breeds. The two-sided *P* value was calculated by the linear mixed regression model. (h) Manhattan plots for another example, highlighting *AKR1B1* in liver, colocalized with protein percent trait (PP4 = 0.96) and located within a strong selection region (*F*_ST_ = 0.17).

Similar with beQTL results, cell-type-specific cseQTL were significantly enriched in the *F*_ST_10 (**Fig. 8a**). Among 23 tested tissue-cell pairs with specific cseQTL, fibroblasts in epididymis, T cells in lymph node, and luminal cells in mammary gland were top cell types under selection between dairy and beef cattle (**Fig. 8b**). A total of 143 selected cseGenes in luminal cells of the mammary gland and T cells of the lymph node were significantly enriched in milk production–related pathways, including the Wnt signaling pathway (FDR = 4 × 10⁻²) and ubiquitin-mediated proteolysis (FDR = 3 × 10⁻⁴) (**Fig. 8b; Supplementary Fig.25a**). Compared to liver beQTL, cseGenes in hepatocytes of liver exhibited more colocalizations with milk production, including fat yield, fat percent, milk yield, net merit, and protein percentage (**Fig. 8c**). For instance, the lead cseQTL (*rs209021141*) of *TMEM249*, a potential causal gene for the milk production in the liver^44^, was colocalized with milk fat percentage (PP4 = 0.90) and resided in a strong selection region (*F*_ST_ = 0.25), which was not observed in the liver beQTL (**Fig. 8d**). The frequency of T allele of *rs209021141* was elevated in the Holstein dairy cattle compared to beef cattle (**Fig. 8e**). In addition, luminal cells of mammary gland harbored four cseGenes that were colocalized with milk yield, net merit, fat percentage, and protein percentage (**Fig. 8c**). For instance, the lead cseQTL (*rs132986937*) of *SLC35B4* in luminal cells was colocalized with protein percentage (PP4 = 0.96) and resided in a strong selection region (*F*_ST_ = 0.17) (**Fig. 8f**). Compared to beef cattle, the T allele of *rs210230814* increased gene expression of *SLC35B4* in luminal cells, and had a higher frequency in the Holstein dairy cattle than beef cattle (**Fig. 8g**). The *SLC45B4* gene regulates glucose metabolism and nutrient transport in luminal cells and may indirectly influence the synthesis and secretion of milk proteins^45^. Similarly, the lead cseQTL (*rs210230814*) of *USP3* in luminal cells of mammary gland, was colocalized with fat percentage (PP4 = 0.79) and was under strong selection (**Supplementary Fig25b**). The G allele of this SNP decreased *USP3* expression and was present at a higher frequency in Holstein cattle than beef cattle (**Supplementary Fig.25c**). Through the role of *USP3* in modulating immune responses, it may influence the immune microenvironment of the mammary gland, thereby impacting lactation^46^. These two genes were also identified as beGenes with strong selection signal and colocalized with milk fat yield and milk protein percentage in mammary gland (**Supplementary Table 12 and 13**). Altogether, these findings highlighted the potential of this CattleCell-GTEx resource in dissecting molecular mechanisms underlying population selection.

**Fig. 8.**
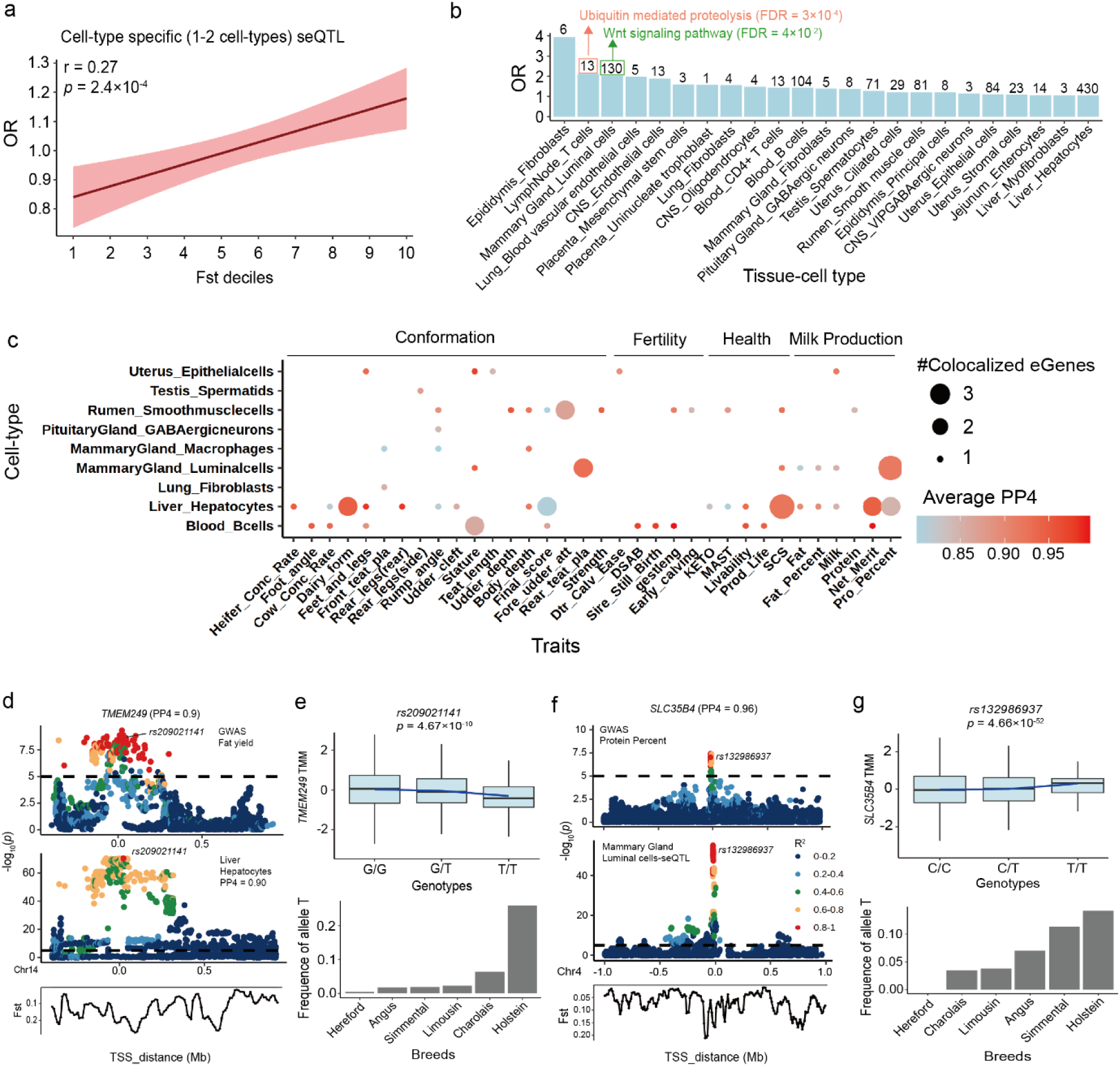
Linking cell-type stratified eQTL (cseQTL) to selection signatures between dairy and beef cattle. (a) Enrichment of cseQTL across 10 *F*_ST_ deciles. The lines were fitted by a linear regression model using the geom_smooth function from ggplot2 (v3.3.2) in R (v4.0.2). The *P* values were obtained by Pearson’s correlation (*r*) test. (b) The number of cseGenes under strong selection (top 10% *F*_ST_) across 23 tissue–cell pairs. Bar height indicates gene counts, and selected KEGG pathways related to milk production are labeled. (c) The bubble plot comparing colocalized and selected cell-type-specific cseGenes across traits. Dot size represents the number of cell-type-specific cseGenes with both strong selection signal and colocalization per trait ,while the dot color indicates the average colocalization posterior probability (PP4). (d) Manhattan plots show an example of a selected and colocalized cseGene (*TMEM249*) in liver hepatocytes, associated with net merit (PP4 = 0.90). The lead variant *rs209021141* lies within a high *F*_ST_ region (*F*_ST_ = 0.25). (e) The boxplot shows expression of *TMEM249* in liver hepatocytes across three genotypes (G/G, G/T, T/T) of *rs209021141*, while the barplot indicates the allele T frequency distribution across five cattle breeds. The two-sided *P* value was calculated by the linear mixed regression *cis*-eQTL model. (f) Manhattan plots highlight *SLC35B4* as a selected and colocalized cseGene in luminal cells of the mammary gland, associated with milk protein percent trait (PP4 = 0.96). The causal variant *rs132986937* is located in a strong selection region (*F*_ST_ = 0.17). (g) The boxplot shows allele-specific expression of *SLC35B4* in luminal cells of mammary gland across three genotypes (C/C, C/T, T/T) of *rs132986937*, while the barplot indicates the allele T frequency distribution across five cattle breeds. The two-sided *P* value was calculated by the linear mixed regression *cis*-eQTL model.

## Discussions

In summary, we presented the CattleCell-GTEx resource—an invaluable and comprehensive catalog of genetic regulatory effects on gene expression across 81 cell types in 22 bovine tissues. This resource included 18,557 beGenes, 168 fQTL, 16,538 cseGenes, 16,341 cell-type ieGenes, and 15,012 cell-state ieGenes. More importantly, this resource offers an in-depth understanding of the genetic regulatory effects on the bovine transcriptome at the cellular level and their impacts on complex traits and population selection in cattle. We demonstrated that different types of eQTL mapping identified unique sets of regulatory variants with distinct tissue and cell type sharing patterns. On average, approximately 75% of GWAS loci from 44 complex traits examined in cattle could be colocalized by at least one of eQTL being detected, among which 32% were attributed by cell-type specific eQTL only, consistent with the proportion (36%) in the previous human analysis for cell-type eQTL^13^. In addition, by applying this CattleCell-GTEx resource to population selection results, we demonstrated the important roles of regulatory variants in cattle selective breeding, which was also reported in humans^47,48^.

Although the current CattleCell-GTEx resource provided novel insights into genetic control of gene expression at cell type level, we noticed several limitations in this resource, and gave perspectives on the future development of CattleCell-GTEx project. Firstly, the limitation was inherent in cell-type deconvolution of bulk RNA-seq samples, including its weak performance on highly correlated cell types/states and rare cell types^49,50^. Secondly, we only investigated common SNPs that could be imputed from RNA-seq data accurately. Thirdly, the prioritized variants, genes, cell types, and tissues for complex traits and selection cannot be considered as ‘causal’ directly, due to the LD among nearby genomic variants, gene expression correlations, and similarities of cell types^51^. In the future, it will be good to directly generate single-cell RNA-seq data at population scale, coupled with rich metadata records and deep whole-genome sequences. This will then allow us to explore the impacts of multiple types of genomic variants (e.g., structure variants, rare variants, and somatic mutations) on gene expression in both common and rare cell types^52,53^. It will be also important to functionally validate the prioritized regulatory variants and genes using either *in vivo* or *in vitro* systems *via* massively parallel report assays and CRISPER-based editing technologies^54,55^.

## Acknowledgments

1. L. F. was supported by Agriculture and Food Research Initiative Competitive grants nos. 2022-67015-36215 from the USDA National Institute of Food and Agriculture, and seed-funding from CellFood Hub (AUFF). G. E. L. was supported in part by the Agriculture and Food Research Initiative Competitive grant numbers 2019-67015-29321 and 2021-67015-33409 from the USDA National Institute of Food and Agriculture (NIFA). D. S. and B. H. supported by the National Key R&D Program of China (2021YFF1000700, 2022YFF1000103, 2024YFF1000100); the National Natural Science Foundation of China (32372836). B. L. acknowledges funding from the UK Biotechnology and Biological Sciences Research Council (BBSRC) with grant BB/X009505/1. We acknowledge the support of the High-Performance Computing Platform of China Agricultural University (Beijing) and the Xihe High-Performance Computing Platform of the National Research Facility for Phenotypic and Genotypic Analysis of Model Animals (Beijing).

## Author contribution statement

L.Fang and H.Li. conceived and designed the project. H.Li, H.Zhang and Z.Wei performed bioinformatic analyses of RNA-Seq data analysis. P.Zhao conducted whole-genome sequence data analysis. H.Li, Q.Zhang, S.Zhu, T.Shi, B.Han, W.Zheng, Y.Hou and F.Wang performed multi-omics and single-cell RNA-Seq data analysis. H.Li conducted eQTL mapping. H.Li, C.Maltecca and J.Wang performed GWAS integrative analysis. H.Li, L.Yang, S.Akter, N.Bhowmik, Y.Ma, and R.Baldwin performed single cell spatial RNA-seq data analysis. H.Li and P.Zhao performed selection region analysis. L.Fang, H.Li, G.Sahana, Z.Cai, G.Larson, J.Grady, D.MacHugh, F.Laurant, M.Gong, X.Zhu, Q.Lin, Y.Xi, D.Zhu, J.Teng, D.Guan, Y,Wang, X.Lu, M.Shan, C.Li, L.Ma, B.Li, B.An and J.Ren contributed to the critical interpretation of analytical results before and during manuscript preparation. Y.Wang, J.Sheng and M.Wang built the CattleCell-GTEx web portal. L.Fang., G.Liu., D.Sun., H.Sun., Y.Jiang and J.Jiang. contributed to the data and computational resources. H.Li and L.Fang drafted the manuscript. All authors read, edited, and approved the final manuscript.

## Online Methods

### Ethics

All animal procedures and experimental protocols were approved by the Institutional Animal Care and Use Committee (IACUC) of the Beltsville Agricultural Research Center (BARC), under protocol number 16-019 (approved on August 10, 2016).

### Data collection and processing

We collected a total of 26,467 bulk RNA-seq data in 46 tissues and single-cell RNA-seq data of 1,793,854 cells in 59 tissues from CattleGTEx (v1)^9^ and Cattle Cell Atlas^19^ projects, respectively. Of them, 22 tissues were shared in both bulk and single cell data and used for subsequent analyses.

#### Bulk RNA-seq data processing

For bulk RNA-seq data, briefly, we first trimmed adaptors and discarded reads with poor quality using fastp^56^ (v.0.39) with parameters of “--trim_front 3 --trim_tail 3 --length_required 36 --cut_right -- cut_window_size 4 --cut_mean_quality 15”. We then aligned clean reads to the *Bos taurus* ARS-UCD1.2 (v.110) cattle reference genome using STAR^57^ (v2.7.0) with parameters as below: --chimSegmentMin 10 --outFilterMismatchNmax 3 --twopassMode Basic. We kept 12,241 samples with more than 10M clean reads and uniquely mapping rates ≥ 60% for subsequent analysis. We extracted the raw read counts of 21,385 Ensembl (*Bos taurus* ARS-UCD1.2) protein coding genes (PCGs) by featureCounts^58^ (v1.5.2) and obtained their normalized expression (that is, transcripts per million (TPM)) using Stringtie^59^ (v2.1.1). PCR duplicates were removed in bam files produced by STAR^57^ aligner previously using the MarkDuplicates module of Genome Analysis Toolkit^60^ (GATK, v4.3.0.0). Low quality samples with expressed PCGs (TPM > 0.1 and raw read counts > 6) number less than 20% out of all genes were also excluded. After filtering, 8,866 bulk tissue samples were retained for subsequent analysis.

#### Genotype imputation from RNA-seq samples

GATK BaseRecalibrator module and ApplyBQSR module were applied to recalibrate base quality score using the Ensembl dbSNP database (https://grch37.ensembl.org/info/genome/variation/index.html, v.108). After this, all the output bam files from GATK were utilized do the genotype imputation with GLIMPSE2^61^ (v.2.0.0). A total of 28,958,211 SNPs were imputed across 8,866 bulk samples, and 4,317,531 SNPs with MAF > 0.05 and INFO > 0.75 were then retained using BCFtools^62^ (v.1.17).

#### Mapping bias removal

To improve the computational efficiency, WASP software (v.0.3.4) was used to remove mapping bias^63^. After genotype VCF file for all samples were obtained, SNPs in each chromosome are extracted separately using BCFtools^62^ (v.1.17), and the script snp2h5 provided in WASP software was utilized to convert these output VCF files into HDF5 files^63^. The HDF5 files were then used to remove mapping bias for each sample. The script find_intersecting_snps.py was used to extract reads overlapping with SNPs, and STAR (v.2.7.10b) aligner was engaged to remap these reads with exactly the same command^57^. After reads where one or more of the allelic versions of the reads failed to map back to the same location were filtered out using filter_remapped_reads.py, duplicate reads were filtered using the rmdup.py or rmdup_pe.py for single end or pair end RNA-seq data, respectively^63^.

#### Single-cell RNA-seq data processing

For single cell data, we only considered 22 matched tissues with bulk samples. Briefly, the raw scRNA-seq data were aligned to the same cattle reference genome (ARS-UCD1.2 v.110) and subjected to barcode assignment and unique molecular identifier (UMI) counting based on the Cell Ranger^64^ (v.7.0.1) pipeline (10**×** Genomics). For each library, ddqcR^65^ (v.0.1.0) was used to remove low-quality cells. In each cell cluster, cells with n_counts and n_genes values lower than two median absolute deviations (MADs) from the median were filtered out. Additionally, cells with a mitochondrial gene percentage larger than 10% were removed. Doublets removal was performed using DoubletFinder^66^ (v.2.0.3). It first averaged the transcriptional profile of randomly chosen cell pairs to create pseudo-doublets and then predicted doublets according to each real cell’s similarity in gene expression to the pseudo-doublets. The threshold for doublets was set as 20% and other parameters as default. Seurat (v4.0.6)^67^ pipeline was used to perform unsupervised clustering on clean cells. Libraries from the same tissue were merged and underwent normalizing and scaling with default parameters. Harmony (v.0.1.1)^68^ was used to correct for batch effects with the resetting parameters (Lambda = 1, Theta = 0.5). Variable genes were determined using Seurat’s Find-VariableGenes function with default parameters (selection.method = “vst”, nfeatures = 2000). Clusters were identified via the FindClusters function (resolution = 0.5) implemented in Seurat using the top 30 principal components and subsequently visualized using the RunUMAP function (reduction = “harmony”). Artificial annotation was performed on each cell cluster based on the marker genes reported in the Cattle Cell Atlas^19^ project. We removed cell types with cell numbers less than 50 cells. Additionally, within each cell type, cells expressing fewer than 200 genes and genes present in fewer than three cells were filtered out. After this, 999,192 cells from 81 cell types in 22 tissues were retained for subsequent analysis.

### Benchmarking methods for cell component deconvolution

To evaluate the performance of cell-type deconvolution pipelines, we benchmarked seven widely used methods, including three marker-based (i.e., CIBERSORTx^21^ (v.1.0), DeconRNASeq^69^ (v.3.21) and BisqueRNA^70^ (v.1.0.5), three reference-based (i.e., DWLS^20^ (v.0.1.0), MuSiC^22^ (v.1.0.0) and SCDC^71^ (v. 0.0.0.9)), and one reference-free approaches (i.e., CDSeq^72^ (v. 1.0.9)). Using single-cell RNA-seq data from the cerebral cortex, heart, and ileum, we created pseudo-bulk datasets by randomly splitting cells of each type—half for reference and half for generating simulated bulk samples. To reflect the complexity of real bulk samples, we simulated 27,000 pseudo-bulk samples under three different distributions of cell type proportion (i.e., uniform, normal, and bimodal normal) with varying types of cells (e.g., 10 in cortex, 15 in heart, 20 in ileum) as below. We then assessed the deconvolution performances of these approaches using Pearson correlation and root mean squared error (RMSE) between estimated and true cell-type proportions across samples.

#### Pseudo-Bulk Construction

Pseudo-bulk samples were generated using three distribution models:

- Uniform: Cell-type fractions sampled from Uniform(0.01, 0.99), constrained to sum to 1.
- Normal: Fractions sampled from Normal(0, sd) with sd=0.1, 0.5, 0.9.
- Bimodal normal: Mixture of Bimodal(0, sd1) and Bimodal(1, sd2).

Gene expression was first normalized to TPM:

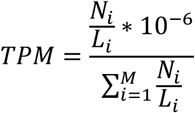

where Ni and Li represent the read count and length of gene i, and M is the total number of genes. Pseudo-bulk expression was computed as:

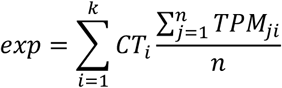

where k is the number of cell types, and CTi is the fraction of cell type i.

#### Marker-based methods

We identified cell-type-specific markers (adj.p < 0.05) with top 50 logFC values per tissue using *Seurat*’s FindMarkers function. These marker genes were used to create signature matrices for deconvolution with CIBERSORT^21^ (v.1.0, SVR-based, 1000 permutations), DeconRNASeq^69^ (v.3.21, quadratic programming, default settings) and BisqueRNA^70^ (v.1.0.5, default settings).

#### Reference-based methods

Using single-cell expression, we constructed *SingleCellExperiment* objects for each tissue and applied DWLS^20^ (v.0.1.0), MuSiC^22^ (v.1.0.0) and SCDC^71^ (v. 0.0.0.9), both under default parameters, for deconvolution.

#### Reference-free methods

We applied CDSeq^72^ (v. 1.0.9), a reference-free method based on latent Dirichlet allocation (LDA), to estimate cell-type proportions from bulk RNA-seq data. To improve consistency with other methods, we provided a reference file with averaged gene expression per cell type, and all other parameters were set to default.

### Cell components deconvolution across bulk samples

To ensure data quality, we applied the following filtering steps to both bulk RNA-seq data and single-cell references. In bulk data, only genes expressed in both datasets were retained and expression levels were converted to TPM. For single-cell references, cell types with fewer than 50 cells were excluded. Within each cell type, cells expressing fewer than 200 genes and genes detected in fewer than three cells were also removed. We then used DWLS to deconvolute cell type fractions of all the bulk RNA-seq samples, where cleaned bulk and single cell reference expression matrices were provided as input, with default parameters.

### Cell-state deconvolution across bulk samples

To construct a cell-state reference, we applied trajectory analysis using Slingshot^73^ (v.2.6.0) to estimate pseudotime for each cell within a given cell type. For cell types with multiple trajectories, only the one containing the most cells was retained and divided into 10 distinct cell-state bins. We then used the MeDuSA^16^ (v.1.0) to estimate cell-state compositions across bulk samples via the MeDuSA_obj function, setting resolution = 10. MeDuSA^16^employs a linear mixed model (LMM) framework, treating the focal state (a single cell or the mean of multiple cells) as a fixed effect, while modeling the remaining cells within the same cell type as random effects to account for intra-type correlation.

### Prediction of cell-type-specific gene expression across bulk samples

Based on the deconvoluted cell-type fractions, we estimated gene expression for each cell type in bulk samples using bMIND^23^ (v.0.3.3) with default parameters. To assess the performance of bMIND, we first tested it on pseudo-bulk data. Specifically, half of the cells in each cell type were used to build a reference expression profile, while the other half formed simulation datasets. Reference profiles were computed as averaged gene expression across cells of the same type. Ten simulated pseudo-bulk samples were generated using Simbu^74^ (v.1.1.5) with default parameters, and cell-type fractions were estimated using DWLS^20^. Gene expression was inferred using bMIND with default parameters. Pearson’s correlation between the inferred and known expression levels was calculated to evaluate the prediction accuracy. Finally, bMIND was applied to real bulk samples across 22 tissues to estimate cell-type-specific gene expression across 49 common cell types (the estimated cell fraction > 5%).

### GWAS analysis of cell fraction

GWAS analysis was performed between genotypes and estimated cell-type fraction in 297 tissue-cell type pairs using the linear mixed model, implemented in Omiga^75^ (v.1.0.3) with the option of “--mode gwas”. Independent SNPs were identified using the GCTA-cojo function in GCTA^76^ (v1.94.1), based on SNPs of each corresponding tissue. We considered independent SNPs with *p*-values less than 5 × 10⁻^8^ as genome-wide significant, and defined cell fraction QTL (fQTL) regions by extracting all SNPs located within ±1 Mb of each independent SNP^77^.

### Association of cell-type fraction QTL (fQTL) with complex traits

We used two thresholds to define the significant association of 168 fQTL with 44 complex traits. Firstly, trait GWAS loci matched with lead SNP in each fQTL were extracted across 44 traits and lead SNP with p-value less than 3×10^-4^ (0.05/168) were defined as candidate associated SNPs between fQTL and complex traits. Next, we performed a large-scale association lookup using genome-wide association summary statistics for each trait to account for potential random effects. In each permutation, 168 SNPs were randomly sampled from the genome-wide set (excluding the candidate SNPs), and the number of SNPs with *p*-values below 3×10^-4^ was recorded. This process was repeated 1,000 times to generate a null distribution of significant SNP counts under random selection. We then compared the observed number of significant candidate SNPs to this null distribution to calculate an empirical *p*-value. An empirical significance threshold was also derived as the 95th percentile of the permutation-based distribution, providing a trait-specific threshold to determine whether the candidate SNPs showed more significant associations than expected by chance. All permutation analyses were conducted using custom R scripts and available to be downloaded (https://github.com/FarmGTEx/CattleCell-GTEx-Pipeline-v0).

### eQTL mapping

For eQTL mapping, we only considered imputed SNPs with INFO > 0.75, MAF≥0.05, and minor allele count≥6, resulting in 4,317,531 SNPs. PCGs with TPM < 0.1 and/or raw read counts < 6 in more than 80% of samples were also removed. Clean gene expression matrix of both bulk and cell-type specific RNA-seq data were normalized to Trimmed Mean of M-values (TMM) values, implemented in edgeR, followed by inverse normal transformation of the TMM. Genotype and gene expression principle components (PCs) were computed using SNPRelate^78^ (v.1.32.2) and PCAforQTL^79^ (v.0.1.0) respectively. We performed *cis*-eQTL mapping using a linear mixed regression model, implemented in Omiga^75^ (v.1.0.3). We defined the *cis*-window of PCG as ±1 Mb of TSS and obtained the nominal *P* values of *cis*-eQTL with the option of “--mode cis”, while selecting the suitable PCs as covariance using the optional of “--dprop-pc-covar 0.001”. Covariances with high co-linear were removed with the option of “--rm-collinear-covar 0.95”. We then employed two layers of multiple testing corrections using Omiga. In the first layer, we applied an adaptive permutation approach to calculate the empirical *P*g values of variants within each gene and obtained the permutation *P*g value of the lead variant for each gene. In the second layer, we conducted the clipper correction for the nominal *P*g values and computed the FDR of lead variants across all tested genes using the option of “--multiple-testing clipper”. The genes with nominal *P*g values less than the permutation *P*g value and FDR less than 0.05 were defined as the genome-wide significant eGenes^75^.

### ieQTL mapping

For cell type or cell state fraction, we only considered cell types with fraction more than 5% and excluded samples that were more than ±3 standard deviations (SD) from the mean^80^. We performed cis-interaction eQTL mapping using estimated cell-type/cell-state fraction as interaction factors with the option of “--mode cis_interaction” i n tested eGenes, implemented in Omiga^75^. Other parameters kept the same with eQTL mapping part. The gene with nominal *P*g*i values less than the permutation *P*g*i value and FDR less than 0.05 were defined as the genome-wide significant ieGenes^75^.

### Fine mapping of eQTL

We first used the stepwise regression procedure for mapping conditionally independent eQTL, as used in other GTEx studies^9,11,12,81^. This analysis was done by using the Omiga with the “–mode cis_independent” option, same to independent test implemented in TensorQTL^82^. The conditionally independent eQTL mapping was based on the nominal associations mentioned above and ranked variants. Secondly, we fine-mapped putative causal variants for gene expression by using the “Sum of Single Effects” (SuSiE) model^83^ (v.1.0). We calculated LD correlations between all tested SNPs from the genotype reference panel using Plink^84^ (v.1.9) and then fine-mapped variants using the SuSiE infinitesimal effect model.

### The tissue/cell type-sharing patterns of eQTL

To understand the shared or specific genetic regulatory mechanisms under tissue and cell-type levels, we performed a meta-analysis of eQTL using MashR^85^ (v0.2.6). We only considered the *z* scores (slope/slope_se) from Omiga of the top *cis*-eQTL. We obtained the estimated effect sizes (that is, posterior means) and the corresponding significance levels (that is, local false sign rate (LFSR)) from the mash function. We defined an eQTL with LFSR < 0.05 as active in a given tissue or cell type. To estimate the pairwise tissue/cell-type similarity with regard to genetic regulation of gene expression, we calculated the pairwise Spearman’s correlation of effect size estimates of *cis*-eQTL between any tissue or cell-type pairs, focusing on SNPs with LFSR < 0.05 in at least one tissue/cell-type^11^.

### GWAS summary statistics of complex traits

To investigate the regulatory mechanisms underlying complex traits in cattle, we systematically integrated identified eQTL with GWAS summary statistics for 44 complex traits in Holstein cattle^86^, including 22 body conformation, 6 milk production, 8 reproduction, and 8 health traits. Independent SNPs were identified for each trait using the *GCTA-cojo* function in GCTA^76^, based on a Holstein reference genomic panel. For colocalization analysis, independent SNPs with *p*-values less than 1 × 10⁻⁵ were then selected as suggestive significant, and corresponding QTL regions were defined by extracting all SNPs located within ±1 Mb of each lead SNP.

### *Cis*-eQTL-GWAS colocalization

To identify shared genetic variants between GWAS variants and eQTL/ieQTL, we conducted a colocalization analysis between each eGene/ieGene and overlapped GWAS QTL regions using the *coloc.abf* function in the *coloc* package^87^ (v.5.2.3), which is an approximate Bayes factor colocalization analysis for detecting significant genetic variants shared by GWAS variants and eQTL/ieQTL. The package computed posterior probabilities for (1) no association with either GWAS variants and eQTL/ieQTL (*H*0); (2) association only with GWAS variants (*H*1); (3) association only with eQTL/ieQTL (*H*2); (4) association with both GWAS variants and eQTL/ieQTL but two independent signals (*H*3); and (5) association with both molecular phenotype and shared signals (*H*4). Moreover, we calculated the LD of two lead SNPs for a pair of GWAS variants and eQTL/ieQTL using PLINK^84^ (v.1.9).

### TWAS of complex traits

We conducted a transcriptome-wide association study (TWAS) with S-PrediXcan^88^ function, implemented in the MetaXcan^89^ (v.0.6.11) family. In brief, we trained the nested cross-validated elastic net models with gene expression and corresponding SNPs within the 1 Mb *cis*-window of gene expression in all 22 tissues and 72 tissue-cell type pairs. The predictive models with cross-validated correlation ρ > 0.1 and prediction performance *P* < 0.05 were selected for subsequent analyses. Using the S-PrediXcan tool and trained models, we predicted gene–trait associations at the single-tissue and cell-type level, and defined significant associations using FDR less than 0.05.

### Detection of trait-relevant cell types via enrichment analysis

The Scpagwas^90^ (v.1.3.0) software package was employed to perform the enrichment analysis between cell types and complex traits. Briefly, scPagwas uses a polygenic regression model to prioritize a set of trait-relevant genes and uncover trait-relevant cell subpopulations by incorporating pathway activity transformed scRNA-seq data with GWAS summary data. To enhance the comprehensiveness of our results, 319 human KEGG pathways downloaded from the KEGG database^91^ were used after eliminating duplicates and converting homologous genes. The Boot_evaluate function was employed to identify the significant trait-relevant relevant cell types and calculate trait-relevant scores. The scGet_PCC function was used to prioritize the top trait-relevant genes by ranking the Pearson correlation coefficient (PCC). Genes with the top 50 PCC values were defined as trait-relevant genes in each cell type. In addition, the scPagwas_perform_score function was applied to perform pathway activity analysis and define the significance of active pathways in each cell type based on the singular value decomposition (SVD) method. Enrichment analysis between trait-relevant genes and active pathway genes was performed based on Fisher’s test and Chi-square test within each cell type using all expressed genes as background.

### Spatial transcriptome data processing

#### Tissue collection and processing

Prefrontal cortex tissue was collected from a Holstein cow as part of the ongoing FarmGTEx-Cattle initiative to establish a comprehensive tissue bank for gene expression studies. Following euthanasia in accordance with approved BARC protocols, brain tissues were rapidly dissected. The prefrontal cortex was carefully isolated, rinsed with cold sterile phosphate-buffered saline (PBS) to remove residual blood, and embedded in optimal cutting temperature (OCT) compound on dry ice. Samples were immediately snap-frozen in liquid nitrogen and stored at −80°C until further processing.

#### RNA extraction, library preparation, and sequencing

Total RNA was extracted using the RNeasy Plus Mini Kit (Qiagen, Germany) following the manufacturer’s instructions. RNA integrity was assessed using the Agilent 2100 Bioanalyzer (Agilent Technologies, USA), and only samples with RNA Integrity Number (RIN) ≥ 8.0 were used for library construction. mRNA-seq libraries were prepared using the Illumina TruSeq Stranded mRNA Library Prep Kit (Illumina, San Diego, CA) according to the manufacturer’s protocols. Library quality and concentration were evaluated using the Qubit 4.0 Fluorometer and the Agilent Bioanalyzer. Libraries were sequenced on an Illumina NovaSeq 6000 platform using paired-end 150 bp reads.

#### Cell-type and brain region annotation

Expression matrix and spatial information were extracted and saved as h5ad file with bin 50. Law quality cells with expressed genes less than 3, read counts less than 20 and gene counts by read counts more than 4000 were removed. Additionally, cells with a mitochondrial genes percentage higher than 5% were also removed. The clustering process was completed based on Stereopy^92^ (v.1.6.0). Firstly, high variable genes (HVGs) were determined using Stereopy’s *highly_variable_genes* function with default parameters (n_top_genes = 2000). Clusters were identified via the *leiden* method (resolution = 1.3) implemented in Stereopy using the top 30 principal components and subsequently visualized using the *cluster_scatter* function (res_key=’leiden’). Single cell data of cerebral cortex from CattleCA^19^ dataset and the expression matrix of 13 human brain regions (https://www.proteinatlas.org/humanproteome/brain/data) were utilized to build reference for annotation. Top 50 genes of log2FC in each cell type or brain regions were extracted into the reference expression matrix. Annotation was completed based on SingleR^93^ (v.2.0.0) software with the default parameters. The annotated data was saved as h5ad file to perform the following enrichment analysis.

### Enrichment between spatial cell types and GWAS summary data

gsMap^94^ (v.1.73.5) was utilized to define the association of cell types with complex traits in the spatial level. Firstly, gsMap used a graph neural network (GNN) to identify homogeneous spots for each focal spot in terms of both gene expression patterns and spatial positions using the *run_find_latent_representations* function. Next, the gene specificity scores (GSSs) of each spot were computed by aggregating information from these homogeneous spots, representing the relative rank of the expression level of each gene in a spot, using the *run_latent_to_gene* funciton. Second, we assigned GSS to SNPs and computed the stratified LD score using the *run_generate_ldscore* dunction. Finally, the *run_spatial_ldsc* function was utilized to calculate spatial LDSC to associate spots with a trait.

### Function enrichment analysis of gene lists

Gene Ontology (GO) and Kyoto Encyclopedia of Genes and Genomes (KEGG) analyses were performed using the clusterProfiler^95^ (v.4.0). The GO terms and KEGG pathways of selected genes were enriched in the org.Bt.eg.db and KEGG-bta databases, using the erichGO and enrichKEGG functions, respectively, with a threshold parameter of “pvalueCutoff = 0.05”. We considered a GO term or a KEGG pathway significant if it had FDR < 0.05.

### Selective sweep between dairy and beef cattle

Selective sweeps across cattle populations were identified between 651 dairy (Holstein) and 552 beef (326 Simmental, 64 Angus, 60 Charolais, 58 Limousin and 44 Hereford) cattle using genome-wide SNPs (**Supplementary Table 10**). We analyzed genetic differentiation (*F*st) using the vcftools^96^ (v.0.1.16) tool, employing a sliding window of 30 Kb with a step size of 10 Kb.

### Enrichment of eQTL in selection signatures

The windows were divided into deciles based on their Fst values, forming ten groups (*F*st1–*F*st10) with increasing selection intensity. Enrichment analysis was then performed between each group and the eQTL from credible sets derived through fine mapping. Odd ratio (OR) was calculated to evaluate enrichment levels between selected regions and eQTL in with the following formula:

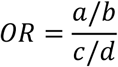

In which a represents the numbers of eQTL located in genomic regions of each *F*st groups, b represents the number of eQTL, c represents the number of SNPs located in genomic regions of each *F*st groups, d represents the number of all SNPs. The frequency of alleles in each SNP was calculated using the option of “--freq” in Plink.

### Statistics and reproducibility

No statistical method was used to predetermine the sample size. The details of data exclusions for each specific analysis are available in the <u>Methods</u> section. For all the boxplots, the horizontal lines inside the boxes show the medians. Box bounds show the lower quartile (*Q*1, the 25th percentile) and the upper quartile (*Q*3, the 75th percentile). Whiskers are minima (*Q*1 − 1.5× IQR) and maxima (*Q*3 + 1.5× IQR), where IQR is the interquartile range (*Q*3–*Q*1). Outliers are shown in the boxplots unless otherwise stated. The experiments were not randomized, as all the datasets are publicly available from observational studies. The investigators were not blinded to allocation during experiments and outcome assessment, as the data were not from controlled randomized studies.

## Data availability

All raw data analyzed in this study are publicly available for download without restrictions from SRA (https://www.ncbi.nlm.nih.gov/sra/). The ARS-UCD1.2 (v.110) cattle reference genome is available at Ensembl (https://www.ensembl.org). Details of RNA-Seq samples, eQTL summary results, selection analysis results, and ATAC peaks can be found in Supplementary Tables, respectively. All processed data and the full summary statistics of eQTL mapping and genotype imputation reference panel are available at https://cattlecellgtex.farmgtex.org/.

## Code availability

All the computational scripts and codes for RNA-Seq and WGS analyses, as well as the respective quality control, deconvolution, molecular phenotype normalization, eQTL mapping, functional enrichment, colocalization, TWAS and selection signal are available at the FarmGTEx GitHub website (https://github.com/FarmGTEx/CattleCell-GTEx).

## References

1. Bruford, M.W., Bradley, D.G. & Luikart, G. DNA markers reveal the complexity of livestock domestication. Nature Reviews Genetics 4, 900–910 (2003).

2. Smith, J. et al. Beyond milk, meat, and eggs: Role of livestock in food and nutrition security. Animal Frontiers 3, 6–13 (2013).

3. Tona, G.O. Impact of Beef and Milk Sourced from Cattle Production on Global Food Security. in Bovine Science - Challenges and Advances (ed. Abubakar, M.) (IntechOpen, Rijeka, 2021).

4. Schultz, B., Serão, N. & Ross, J.W. Chapter 23 - Genetic improvement of livestock, from conventional breeding to biotechnological approaches. in Animal Agriculture (eds. Bazer, F.W., Lamb, G.C. & Wu, G.) 393–405 (Academic Press, 2020).

5. Xia, X. et al. Global dispersal and adaptive evolution of domestic cattle: a genomic perspective. Stress Biology 3, 8 (2023).

6. Chen, N. et al. Global genetic diversity, introgression, and evolutionary adaptation of indicine cattle revealed by whole genome sequencing. Nature Communications 14, 7803 (2023).

7. Ritter, C.A.-O., Beaver, A.A.-O. & von Keyserlingk, M.A.-O. The complex relationship between welfare and reproduction in cattle.

8. Laborde, D., Mamun, A., Martin, W., Piñeiro, V. & Vos, R. Agricultural subsidies and global greenhouse gas emissions. Nature Communications 12, 2601 (2021).

9. Liu, S. et al. A multi-tissue atlas of regulatory variants in cattle. Nature Genetics 54, 1438–1447 (2022).

10. Clark, E.L. et al. From FAANG to fork: application of highly annotated genomes to improve farmed animal production. Genome Biology 21, 285 (2020).

11. Teng, J. et al. A compendium of genetic regulatory effects across pig tissues. Nature Genetics 56, 112–123 (2024).

12. Guan, D. et al. Genetic regulation of gene expression across multiple tissues in chickens. Nature Genetics 57, 1298–1308 (2025).

13. Kim-Hellmuth, S. et al. Cell type–specific genetic regulation of gene expression across human tissues. Science 369, eaaz8528 (2020).

14. Kasela, S. et al. Interaction molecular QTL mapping discovers cellular and environmental modifiers of genetic regulatory effects.

15. Nathan, A. et al. Single-cell eQTL models reveal dynamic T cell state dependence of disease loci. Nature 606, 120–128 (2022).

16. Song, L., Sun, X., Qi, T. & Yang, J. Mixed model-based deconvolution of cell-state abundances (MeDuSA) along a one-dimensional trajectory. Nature Computational Science 3, 630–643 (2023).

17. Donovan, M.K.R., D’Antonio-Chronowska, A., D’Antonio, M.A.-O. & Frazer, K.A.-O. Cellular deconvolution of GTEx tissues powers discovery of disease and cell-type associated regulatory variants.

18. Sheng, X.A.-O. et al. Mapping the genetic architecture of human traits to cell types in the kidney identifies mechanisms of disease and potential treatments.

19. Fang, L., et al. Cattle Cell Atlas: a multi-tissue single cell expression repository for advanced bovine genomics and comparative biology. (2024).

20. Tsoucas, D. et al. Accurate estimation of cell-type composition from gene expression data. Nature Communications 10, 2975 (2019).

21. Steen, C.B., Liu, C.L., Alizadeh, A.A. & Newman, A.M. Profiling Cell Type Abundance and Expression in Bulk Tissues with CIBERSORTx.

22. Wang, X., Park, J., Susztak, K., Zhang, N.R. & Li, M. Bulk tissue cell type deconvolution with multi-subject single-cell expression reference. Nature Communications 10, 380 (2019).

23. Wang, J.A.-O., Roeder, K. & Devlin, B. Bayesian estimation of cell type-specific gene expression with prior derived from single-cell data.

24. Terman, B.I. et al. Identification of the KDR tyrosine kinase as a receptor for vascular endothelial cell growth factor. Biochemical and Biophysical Research Communications 187, 1579–1586 (1992).

25. Kern, C. et al. Functional annotations of three domestic animal genomes provide vital resources for comparative and agricultural research. Nature Communications 12, 1821 (2021).

26. Cassandri, M. et al. Zinc-finger proteins in health and disease. Cell Death Discovery 3, 17071 (2017).

27. Hand, J.M. et al. Discovery of a novel oocyte-specific Krüppel-associated box domain-containing zinc finger protein required for early embryogenesis in cattle. Mechanisms of Development 144, 103–112 (2017).

28. Di Giulio, V. et al. The dual nature of KLHL proteins: From cellular regulators to disease drivers. European Journal of Cell Biology 104, 151483 (2025).

29. Mishra, S.K. et al. Emerging roles for human glycolipid transfer protein superfamily members in the regulation of autophagy, inflammation, and cell death. Progress in Lipid Research 78, 101031 (2020).

30. Schulz, V. et al. Functional spectrum and specificity of mitochondrial ferredoxins FDX1 and FDX2. Nature Chemical Biology 19, 206–217 (2023).

31. Sun, L., Su, Y., Jiao, A., Wang, X. & Zhang, B. T cells in health and disease. Signal Transduction and Targeted Therapy 8, 235 (2023).

32. Nemazee, D. Mechanisms of central tolerance for B cells. Nature Reviews Immunology 17, 281–294 (2017).

33. Twigger, A.-J. et al. Transcriptional changes in the mammary gland during lactation revealed by single cell sequencing of cells from human milk. Nature Communications 13, 562 (2022).

34. Wellnitz, O. & Bruckmaier, R.M. Invited review: The role of the blood-milk barrier and its manipulation for the efficacy of the mammary immune response and milk production.

35. Yukinaga, H. & Miyamichi, K. Oxytocin and neuroscience of lactation: Insights from the molecular genetic approach. Neuroscience Research 216, 104873 (2025).

36. Smith, M.S. Lactation alters neuropeptide-Y and proopiomelanocortin gene expression in the arcuate nucleus of the rat. Endocrinology 133, 1258–1265 (1993).

37. Brown, M.L. et al. A macrophage receptor for apolipoprotein B48: cloning, expression, and atherosclerosis.

38. Maxman, G., van Marle-Köster, E., Lashmar, S.F. & Visser, C. Selection signatures associated with adaptation in South African Drakensberger, Nguni, and Tuli beef breeds. Tropical Animal Health and Production 57, 13 (2024).

39. Ghoreishifar, S.M. et al. Signatures of selection reveal candidate genes involved in economic traits and cold acclimation in five Swedish cattle breeds.

40. Bertolini, F. et al. Signatures of selection are present in the genome of two close autochthonous cattle breeds raised in the North of Italy and mainly distinguished for their coat colours. Journal of Animal Breeding and Genetics 139, 307–319 (2022).

41. Illa, S.K., Mukherjee, S., Nath, S. & Mukherjee, A. Genome-Wide Scanning for Signatures of Selection Revealed the Putative Genomic Regions and Candidate Genes Controlling Milk Composition and Coat Color Traits in Sahiwal Cattle.

42. Hosseinzadeh, S., Rafat, S.A., Javanmard, A. & Fang, L. Identification of candidate genes associated with milk production and mastitis based on transcriptome-wide association study. Animal Genetics 55, 430–439 (2024).

43. Rižner, T.L. Enzymes of the AKR1B and AKR1C Subfamilies and Uterine Diseases.

44. Križanac, A.-M. et al. Sequence-based GWAS in 180,000 German Holstein cattle reveals new candidate variants for milk production traits. Genetics Selection Evolution 57, 3 (2025).

45. Zhang, Y., Zhang, Y., Sun, K., Meng, Z. & Chen, L. The SLC transporter in nutrient and metabolic sensing, regulation, and drug development.

46. Duan, T.A.-O. et al. USP3 plays a critical role in the induction of innate immune tolerance.

47. Rinker, D.C. et al. Neanderthal introgression reintroduced functional ancestral alleles lost in Eurasian populations. Nature Ecology & Evolution 4, 1332–1341 (2020).

48. Taylor, D.J. et al. Sources of gene expression variation in a globally diverse human cohort. Nature 632, 122–130 (2024).

49. Avila Cobos, F., Alquicira-Hernandez, J., Powell, J.E., Mestdagh, P. & De Preter, K. Benchmarking of cell type deconvolution pipelines for transcriptomics data. Nature Communications 11, 5650 (2020).

50. Meng, G., et al. imply: improving cell-type deconvolution accuracy using personalized reference profiles.

51. Mountjoy, E. et al. An open approach to systematically prioritize causal variants and genes at all published human GWAS trait-associated loci. Nature Genetics 53, 1527–1533 (2021).

52. Yazar, S. et al. Single-cell eQTL mapping identifies cell type–specific genetic control of autoimmune disease. Science 376, eabf3041.

53. Natri, H.M. et al. Cell-type-specific and disease-associated expression quantitative trait loci in the human lung. Nature Genetics 56, 595–604 (2024).

54. Tewhey, R. et al. Direct Identification of Hundreds of Expression-Modulating Variants using a Multiplexed Reporter Assay.

55. Freimer, J.A.-O. et al. Systematic discovery and perturbation of regulatory genes in human T cells reveals the architecture of immune networks.

56. Chen, S., Zhou, Y., Chen, Y. & Gu, J. fastp: an ultra-fast all-in-one FASTQ preprocessor.

57. Dobin, A. et al. STAR: ultrafast universal RNA-seq aligner.

58. Liao, Y., Smyth Gk Fau - Shi, W. & Shi, W. featureCounts: an efficient general purpose program for assigning sequence reads to genomic features.

59. Pertea, M. et al. StringTie enables improved reconstruction of a transcriptome from RNA-seq reads.

60. McKenna, A., et al. The Genome Analysis Toolkit: a MapReduce framework for analyzing next-generation DNA sequencing data.

61. Rubinacci, S., Ribeiro, D.M., Hofmeister, R.J. & Delaneau, O. Efficient phasing and imputation of low-coverage sequencing data using large reference panels. Nature Genetics 53, 120–126 (2021).

62. Danecek, P. et al. Twelve years of SAMtools and BCFtools. LID - 10.1093/gigascience/giab008 [doi] LID - giab008.

63. van de Geijn, B., McVicker, G., Gilad, Y. & Pritchard, J.K. WASP: allele-specific software for robust molecular quantitative trait locus discovery. Nature Methods 12, 1061–1063 (2015).

64. Zheng, G.X.Y. et al. Massively parallel digital transcriptional profiling of single cells. Nature Communications 8, 14049 (2017).

65. Subramanian, A., Alperovich, M., Yang, Y. & Li, B. Biology-inspired data-driven quality control for scientific discovery in single-cell transcriptomics. Genome Biology 23, 267 (2022).

66. McGinnis, C.S., Murrow, L.M. & Gartner, Z.J. DoubletFinder: Doublet Detection in Single-Cell RNA Sequencing Data Using Artificial Nearest Neighbors.

67. Hao, Y. et al. Integrated analysis of multimodal single-cell data. Cell 184, 3573–3587 e29 (2021).

68. Korsunsky, I. et al. Fast, sensitive and accurate integration of single-cell data with Harmony. Nat Methods 16, 1289–1296 (2019).

69. Gong, T. & Szustakowski, J.D. DeconRNASeq: a statistical framework for deconvolution of heterogeneous tissue samples based on mRNA-Seq data. Bioinformatics 29, 1083–5 (2013).

70. Jew, B. et al. Accurate estimation of cell composition in bulk expression through robust integration of single-cell information. Nature Communications 11, 1971 (2020).

71. Dong, M. et al. SCDC: bulk gene expression deconvolution by multiple single-cell RNA sequencing references. Briefings in Bioinformatics 22, 416–427 (2021).

72. Kang, K. et al. CDSeq: A novel complete deconvolution method for dissecting heterogeneous samples using gene expression data. PLoS Comput Biol 15, e1007510 (2019).

73. Street, K. et al. Slingshot: cell lineage and pseudotime inference for single-cell transcriptomics.

74. Dietrich, A. et al. SimBu: bias-aware simulation of bulk RNA-seq data with variable cell-type composition. Bioinformatics 38, ii141–ii147 (2022).

75. Teng, J., et al. OmiGA: A Toolkit for Ultra-efficient Molecular Trait Analysis in Complex Populations. bioRxiv, 2024.12.19.629424 (2024).

76. Yang, J., Lee, S.H., Goddard, M.E. & Visscher, P.M. GCTA: A Tool for Genome-wide Complex Trait Analysis. The American Journal of Human Genetics 88, 76–82 (2011).

77. Jannot, A.-S., Ehret, G. & Perneger, T. P < 5 × 10−8 has emerged as a standard of statistical significance for genome-wide association studies. Journal of Clinical Epidemiology 68, 460–465 (2015).

78. Zheng, X. et al. A high-performance computing toolset for relatedness and principal component analysis of SNP data.

79. Zhou, H.J., Li, L., Li, Y., Li, W. & Li, J.J. PCA outperforms popular hidden variable inference methods for molecular QTL mapping. Genome Biology 23, 210 (2022).

80. Kim-Hellmuth, S. et al. Cell type-specific genetic regulation of gene expression across human tissues. Science 369(2020).

81. The GTEx Consortium atlas of genetic regulatory effects across human tissues.

82. Taylor-Weiner, A. et al. Scaling computational genomics to millions of individuals with GPUs. Genome Biology 20, 228 (2019).

83. Wang, G., Sarkar, A., Carbonetto, P. & Stephens, M. A simple new approach to variable selection in regression, with application to genetic fine mapping.

84. Purcell, S., et al. PLINK: a tool set for whole-genome association and population-based linkage analyses.

85. Urbut, S.M., Wang, G., Carbonetto, P. & Stephens, M. Flexible statistical methods for estimating and testing effects in genomic studies with multiple conditions. Nature Genetics 51, 187–195 (2019).

86. Jiang, J. et al. Functional annotation and Bayesian fine-mapping reveals candidate genes for important agronomic traits in Holstein bulls. Communications Biology 2, 212 (2019).

87. Wallace, C. et al. Statistical colocalization of monocyte gene expression and genetic risk variants for type 1 diabetes.

88. Barbeira, A.N. et al. Exploring the phenotypic consequences of tissue specific gene expression variation inferred from GWAS summary statistics. Nature Communications 9, 1825 (2018).

89. Barbeira, A. et al. MetaXcan: Summary Statistics Based Gene-Level Association Method Infers Accurate PrediXcan Results. bioRxiv, 045260 (2016).

90. Ma, Y. et al. Polygenic regression uncovers trait-relevant cellular contexts through pathway activation transformation of single-cell RNA sequencing data. Cell Genomics 3, 100383 (2023).

91. Kanehisa, M., Sato, Y., Kawashima, M., Furumichi, M. & Tanabe, M. KEGG as a reference resource for gene and protein annotation.

92. Fang, S. et al. Stereopy: modeling comparative and spatiotemporal cellular heterogeneity via multi-sample spatial transcriptomics. Nature Communications 16, 3741 (2025).

93. Aran, D. et al. Reference-based analysis of lung single-cell sequencing reveals a transitional profibrotic macrophage. Nature Immunology 20, 163–172 (2019).

94. Song, L., Chen, W., Hou, J., Guo, M. & Yang, J. Spatially resolved mapping of cells associated with human complex traits. Nature 641, 932–941 (2025).

95. Wu, T., et al. clusterProfiler 4.0: A universal enrichment tool for interpreting omics data.

96. Danecek, P. et al. The variant call format and VCFtools.

